# Host recovery after skin barrier disruption is individual-specific and associated with microbial functions

**DOI:** 10.64898/2026.03.25.714117

**Authors:** Aarthi Ravikrishnan, Stephen Wearne, Xiuting Li, Ghayathri Balasundaram, Ahmad Nazri Mohamed Naim, Indrik Wijaya, Miao Qin Tay, Alicia Ann May Yap, Poongkulali Rajarahm, Tutie Nooradilah Binte Alui, Connie Tse Kwan Yi, Wan Ling Tan, Yu Zheng Ong, Chris Ho, Bi Renzhe, Amalina Binte Ebrahim Attia, Zhang Ruochong, Steven Thng, Cecilia Brun, Robin Kurfurst, Carine Nizard, Karl Pays, Malini Olivo, Thomas Larry Dawson, John Common, Yi Shan Lim, Niranjan Nagarajan

## Abstract

The human skin is repeatedly exposed to mechanical and environmental stress, particularly in common skin diseases such as eczema, and yet the determinants of recovery remain poorly understood. Using longitudinal, multimodal profiling of skin physiology, structure (Raman spectroscopy), and microbial communities (shotgun metagenomics), we investigated in a human cohort (n=36 subjects, ×2 sites, ×6 timepoints) how host-microbe interactions could jointly shape recovery. Despite baseline variability in physiological parameters, we established that our protocol enables a defined disruption of the stratum corneum. While recovery trajectories for host attributes were notably consistent across age groups and body sites, individual-specific differences in recovery timelines were observed. To assess the role of the skin microbiome, several key time-dependent changes in microbial species were identified including enrichment of select *Cutibacterium* and *Staphylococcus* species and depletion of *Corynebacterium* and *Malassezia* species. Clustering of microbiome stability profiles across subjects and sites identified 6 distinct groups which associate with varying host-recovery patterns and microbial functions. Finally, joint hazards modelling of recovery timing revealed significant contributions from microbial taxa, functions and stability groups, highlighting the under-appreciated role of host-microbial interactions in response to skin stress and in the recovery process.

## Introduction

Human skin forms a primary physiological barrier that protects the body from external environmental insults, including pathogens, pollutants and ultraviolet radiation^1^. As the first line of defence, the skin is continuously exposed to physical, chemical and biological stressors^2^ which, together with intrinsic aging^3^, drive progressive changes in skin structure and function. The cumulative impact of these exposures over time results in alterations in barrier integrity, elasticity, hydration and immune responses^4, 5^. In addition to chronic exposures, the barrier is frequently disrupted by acute mechanical and inflammatory perturbations, including itch-driven scratching^6, 7^ and inflammatory conditions such as Atopic Dermatitis^8^. These events can transiently damage the epidermal barrier and reshape the local microbial ecosystem^9^, yet the processes through which the skin restores barrier integrity and microbial balance following such perturbations remain poorly understood^10^. Emerging evidence highlights the central role of the skin microbiome in maintaining cutaneous health, with host–microbe interactions within the skin holobiont contributing critically to homeostasis^11, 12^. Together, these observations underscore the need to systematically study the host and microbial factors that modulate skin recovery and resilience following barrier disruption.

To investigate skin recovery under well-defined perturbations, experimental stress models have been established to systematically examine how the skin recovers post environmental and physiological challenges^13–18^. Among these, the tape-stripping model is one of the most widely used^19–23^, which induces a transient and reproducible disruption of the epidermal barrier through the sequential removal of stratum corneum layers, resulting in a measurable increase in transepidermal water loss (TEWL). This approach enables longitudinal assessment of barrier repair alongside coordinated changes in skin biophysical properties and the microbial ecosystem^24–26^. However, previous studies of skin stress and recovery have largely examined isolated physiological parameters^21, 23, 26–28^ and have been conducted predominantly in Western cohorts^27, 29^. Additionally, the complex interplay between skin structure, physiology, and the microbiome requires multimodal and longitudinal approaches to resolve the temporal organization of these processes and to define individual-specific trajectories of skin resilience and repair.

To address these gaps, we conducted a large-scale cohort study designed for a systematic investigation of multimodal skin barrier recovery. Combining extensive skin physiological and functional characterization (including with multi-spectral Raman spectroscopy) and shotgun metagenomic sequencing data (n=432 libraries) over 6 timepoints, we were able to establish that despite consistent skin barrier disruption across age groups and sites, recovery trajectories are highly individual-specific. These are reflected in different skin microbiome stability profiles, which together with specific skin microbial taxa and functions associate strongly with recovery speed and saliency. Together, our results highlight the underappreciated role of host-microbial relationships in defining highly individualized recovery of skin structure and function after acute stress.

## Results

### Establishment of a robust skin stress model in the presence of baseline physiological heterogeneity

In order to interrogate skin recovery dynamics following acute stress, we conducted a longitudinal study involving subjects in two age groups, with young (n=18, 21–31 years) and older (n=18, 55–65 years) women, allowing us to study the impact of age differences on recovery processes (**Figure 1A**; **Supplementary Data File 1**). For each individual, two skin sites in close spatial proximity were sampled (dorsal and ventral forearm), to study shared patterns of recovery despite differences in physical properties and environmental exposure^30–32^. Skin barrier disruption was induced using a standardized tape stripping protocol^18^ until a two-fold increase in transepidermal water loss (TEWL) was achieved on both sites, after which recovery was monitored over a 72-hour period at 6 timepoints (t=0h pre, 0h post, 6h, 24h, 48h and 72h; **Figure 1A**). This time duration was chosen based on a pilot study that showed that most individuals showed substantial recovery in stratum corneum thickness in this period (**Supplementary Figure 1**). At all six timepoints, skin physiological measurements (TEWL, hydration, elasticity), multi-spectral Raman spectroscopy and shotgun metagenomics was used to comprehensively capture changes in skin structure, physiology, and microbiome composition for all individuals (**Figure 1A**; **Methods**).

**Figure 1.**
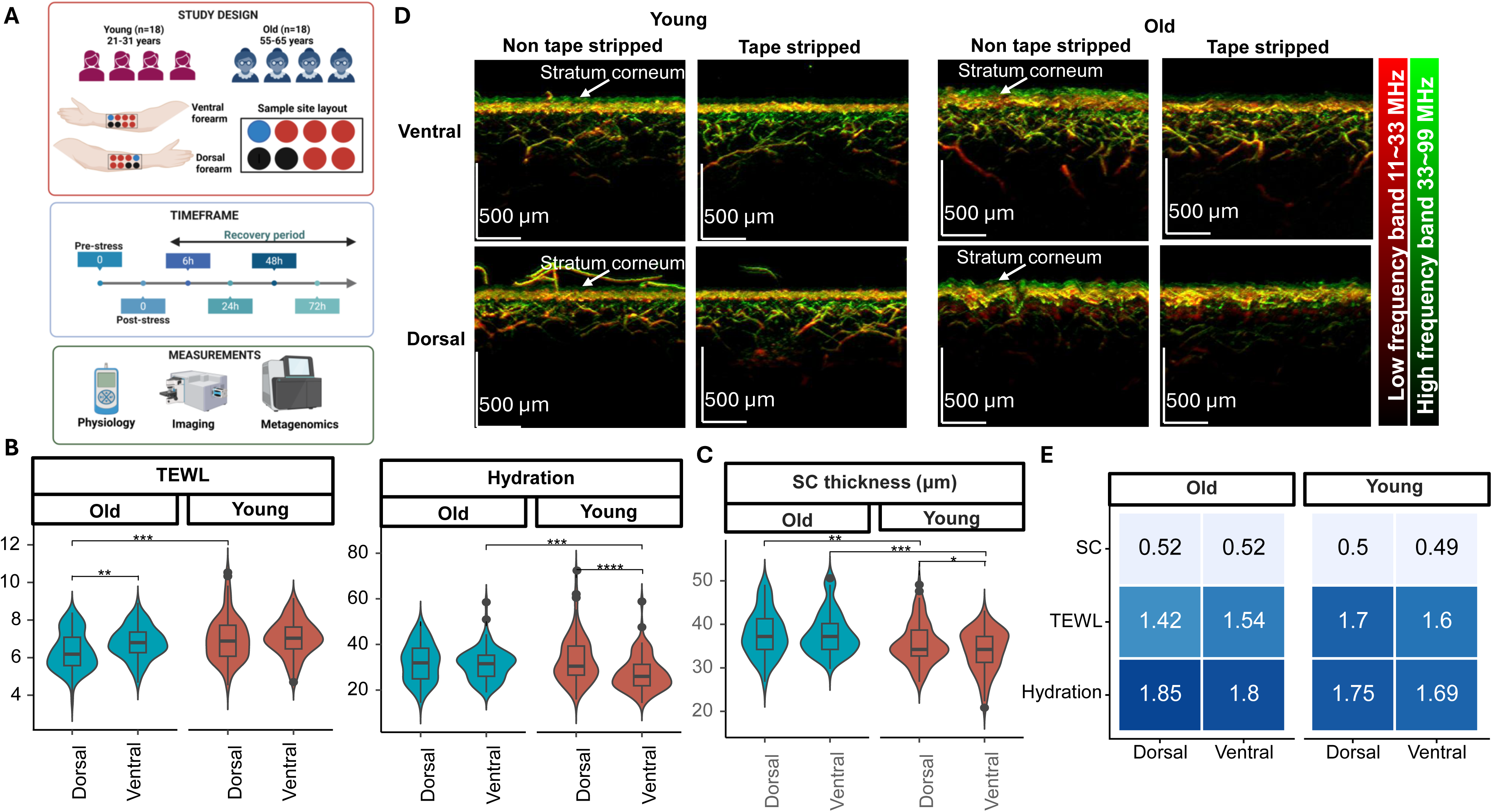
Study design and baseline characteristics of the cohort. (A) Overview of study design showing cohort composition, sampling sites, experimental timeline, and the three sets of measurements collected. The tape-stripped sites (TS; blue and red) and non–tape-stripped (NTS; black) sites are shown, and blue and black sites were used for physiological and imaging measurements at each time point. For microbiome profiling, tape-strip samples were collected from a distinct red site among five marked locations at each time point. (B–C) Violin plots with inset boxplots showing the distribution of (B) transepidermal water loss (TEWL) and hydration, and (C) stratum corneum (SC) thickness (µm) across all timepoints from the NTS site. Paired Wilcoxon tests were used for within-group comparisons, and Wilcoxon rank-sum tests for between-group comparisons. False discovery rate (FDR)–adjusted p-values are denoted as “****” for *p*≤0.0001, “***” for *p*≤0.001, “**” for *p*≤0.01 and “*” for *p*≤0.05. In all boxplots, the center line denotes the median, box limits the upper and lower quartiles, and whiskers the minimum and maximum values. (D) Representative photoacoustic (PA) images pre- and post-tape-stripping in ventral and dorsal sites for younger and older subjects. PA images of both the ventral and dorsal sides were visualized with the LF band (11–33 MHz) in red and the HF band (33–99 MHz) in green at a single wavelength of 532 nm, with larger vascular structures displayed in the LF band and smaller ones in the HF band (Scale bar: 500 µm). (E) Heatmap showing fold-changes in skin parameters calculated as the ratio of tape-stripped and non–tape-stripped sites in younger and older age groups.

Skin physiological measurements revealed significant baseline (t=0h, pre-stress) differences between younger and older individuals in TEWL (higher in younger subjects), hydration (lower in younger subjects) and elasticity (higher in younger subjects; FDR-adjusted Wilcoxon p<0.001; **Figure 1B**, **Supplementary Figure 2**, **Supplementary Data File 2**). Similar statistically significant differences in TEWL and hydration measurements were observed between dorsal and ventral sites, though in all cases the relative differences in median values were <20% (FDR-adjusted Wilcoxon p<0.01; **Figure 1B**). Baseline assessment of skin structure using Raman spectroscopy revealed additional age and site-dependent differences, with significantly higher stratum corneum thickness in older subjects, and younger subjects exhibiting increased stratum corneum thickness in dorsal sites (FDR-adjusted Wilcoxon p<0.05; **Figure 1C**; **Supplementary Data File 2**; **Methods**). Notably, the relative change in median values across groups was small (<10%; **Figure 1C**), though substantial heterogeneity with each group was also observed in many cases (e.g. bimodal distribution for TEWL values in dorsal sites in older subjects; **Figure 1B**, **C**). Interestingly, ventral skin surfaces displayed a smooth and relatively uniform morphology that is not age-dependent, while dorsal skin was visibly wrinkled and uneven in older individuals (**Figure 1D**). Furthermore, quantitative measurements of epidermal and melanin layer thickness, as well as total blood volume exhibited clear differences in younger and older subjects (**Supplementary Figure 2**), highlighting the significant baseline differences across these groups.

To assess the impact of acute stress based on tape stripping, measurements pre- and post-stress (within 20 mins of tape stripping) were compared per individual and site to measure fold change (**Figure 1E**; **Methods**). TEWL measurements exhibited a <2-fold increase across age groups and sites (median range 1.42-1.7), indicating rapid recovery despite the protocol’s effort to calibrate based on this measurement (**Figure 1E**). Hydration measurements were less variable across age groups and sites, but still exhibited <2-fold increase relative to pre-stress values (median range 1.69-1.85; **Figure 1E**). Stratum corneum thickness however exhibited a robust 2-fold reduction across age groups and anatomical sites (median range 0.49-0.52; **Figure 1E**), highlighting the utility of this protocol to deliver a defined skin barrier disruption^33, 34^ across sites despite baseline heterogeneity, thus enabling a robust perturbation framework for downstream study of skin recovery trajectories.

### Recovery of skin physiology follows broadly consistent patterns across age groups and sites

Leveraging data from all 6 timepoints (t=0h pre-stress, t=0h post-stress, t=6, 24, 48 and 72 hours) we next evaluated recovery trajectories across skin physiological parameters (e.g. TEWL, hydration, stratum corneum thickness etc) by comparing data from tape-stripped sites with data from adjacent non-tape-stripped sites that serve as matched controls to compute fold changes (**Figure 2A**; **Supplementary Data File 2**; **Methods**). As expected from our pilot experiment, stratum corneum thickness exhibited recovery by 72 hours across age groups and sites (median fold-change range 0.96-1.01; **Figure 2A**). Hydration displayed a similar recovery trajectory with levels resembling baseline values by 72 hours regardless of age group or site (median fold-change range 1.02-1.15; **Figure 2A**). TEWL values exhibited consistently higher fold-changes for younger subjects across timepoints (FDR-adjusted Wilcoxon p<0.001), but the overall trend of incomplete recovery at 72 hours was shared across age groups and sites (median fold-change>1.1; FDR-adjusted Wilcoxon p<0.01; **Figure 2A**). In contrast, measurements such as elasticity, melanin layer thickness and total blood volume remained stable across timepoints indicating that they were not significantly affected by the mechanical disruption caused by tape stripping in this study (**Supplementary Figure 3**). Integrating all parameters to visualize recovery trajectories (fold-changes) showed that they do not vary significantly across age group or anatomical site, but may be marginally influenced by baseline values for TEWL (PERMANOVA R²=1.7%, p<0.01; **Figure 2B**).

**Figure 2.**
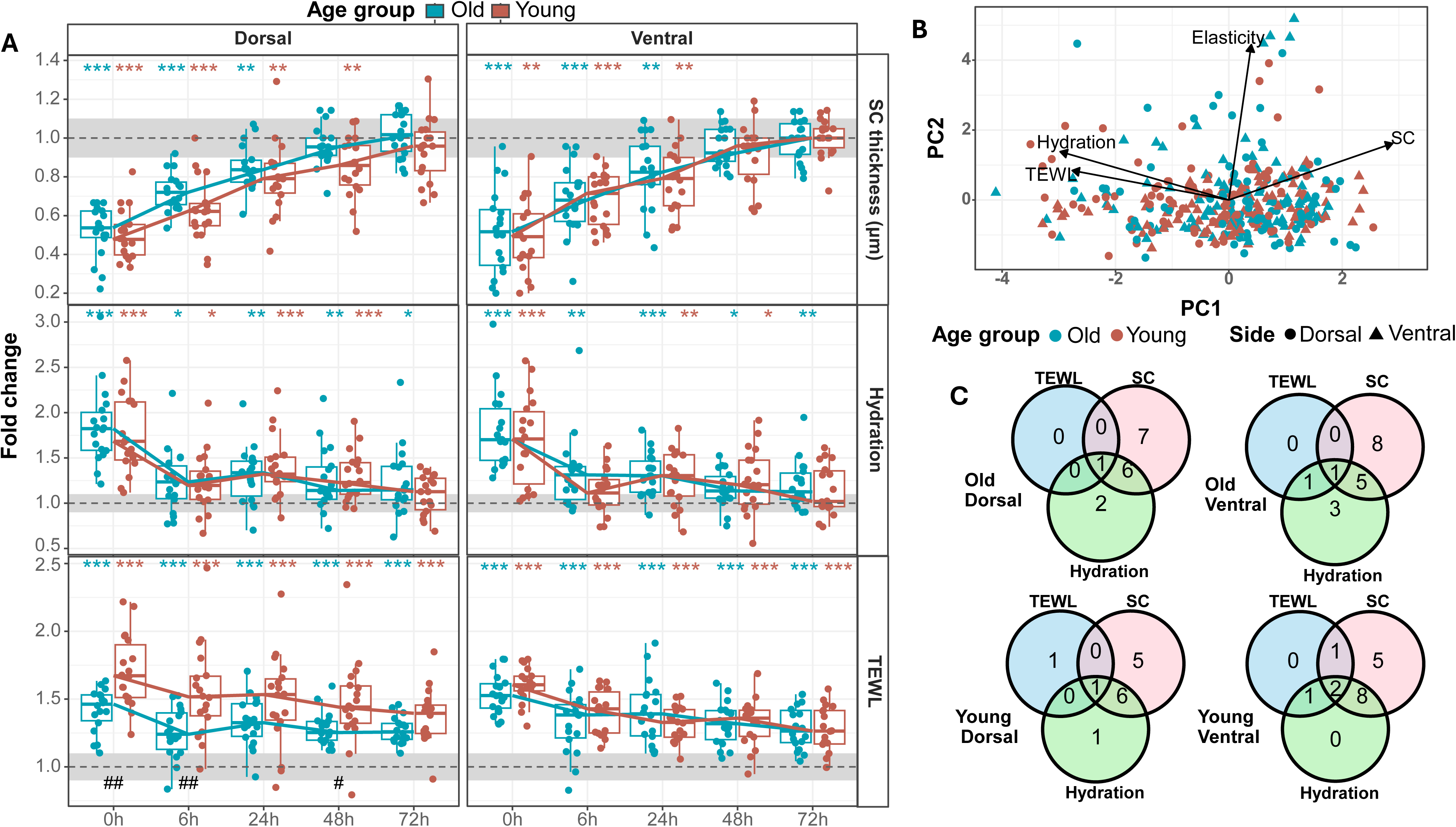
Recovery of skin physiological and structural parameters. (A) Boxplots showing the recovery trajectory measured as the ratio of values from tape-stripped and non-tape-stripped sites (fold change) across timepoints for different parameters: SC thickness, hydration, and TEWL. Within-group comparisons between TS and NTS sites at each timepoint were performed using paired Wilcoxon tests, and between-group (younger vs. older subjects) comparisons using Wilcoxon rank-sum tests on fold-change values. False discovery rate (FDR)–adjusted p-values are indicated as “***” for *p*≤0.001, “**” for *p*≤0.01 and “*” for *p*≤0.05 in within-group tests, and “##” for *p*≤0.01 and “#” for *p*≤0.05 in between-group tests. In all boxplots, center lines denote medians, box limits the upper and lower quartiles, and whiskers the minimum and maximum values. (B) Principal Component Analysis (PCA) plot physiological data from dorsal and ventral sides of younger and older subjects across all timepoints from both non-tape-stripped (NTS) and tape-stripped (TS) sites. Arrows indicate contributions from skin physiological and structural parameters. (C) Venn diagrams showing the number of individuals who achieved recovery in TEWL, hydration, and SC thickness across old dorsal, old ventral, young dorsal and young ventral groups.

To assess recovery at the individual level after 72 hours, we assessed if fold-changes between tape-stripped and non-tape-stripped sites were small (>0.9 or <1.1) similar to baseline variations (t=0h pre-stress; **Methods**). This analysis showed significant consistency across sites and age groups, with shared recovery in stratum corneum thickness and hydration, and non-recovery of TEWL, in many cases (**Figure 2C**). Sites and individuals where all three parameters recovered were not that common but showed patterns of lower baseline stratum corneum values relative to non-recoverers (**Supplementary Figure 4**). Overall, these results highlight the broadly consistent recovery trajectories observed for host physiological parameters across age groups and sites in our tape stripping based acute stress model, despite heterogeneity in baseline measurements.

### Skin microbiome recovers diversity but not composition post stress

Skin tape strips obtained from all subjects and sites, and at all timepoints were used for DNA extraction (n=432) and construction of shotgun metagenomic libraries (n=432, 100% success rate) for short-read sequencing (Illumina, 2×150bp), generating >4 billion reads in total and >9 million reads on average per sample (**Methods**). Read data was uniformly processed together with negative controls to identify and remove potential environmental contaminants, and construct high-quality species-level taxonomic profiles for the skin microbiome (**Supplementary Data File 3**; **Methods**). Baseline skin microbiomes in older individuals exhibited significantly higher alpha diversity metrics across sites (richness, evenness, Shannon & Simpson diversity; **Figure 3A**, **Supplementary Figure 5**), consistent with a depletion of lipophilic species such as *Cutibacterium acnes* (**Supplementary Figure 6**), and an overall shift in community composition (PERMANOVA R²=14%, p<0.001; **Supplementary Figure 6**). Interestingly, despite the age-associated shift, microbiome profiles were highly consistent between dorsal and ventral forearm sites for an individual (Pearson ρ>0.85) and did not carry any significant differentially abundant taxa (**Supplementary Figure 7**, **Supplementary Data File 4**).

**Figure 3.**
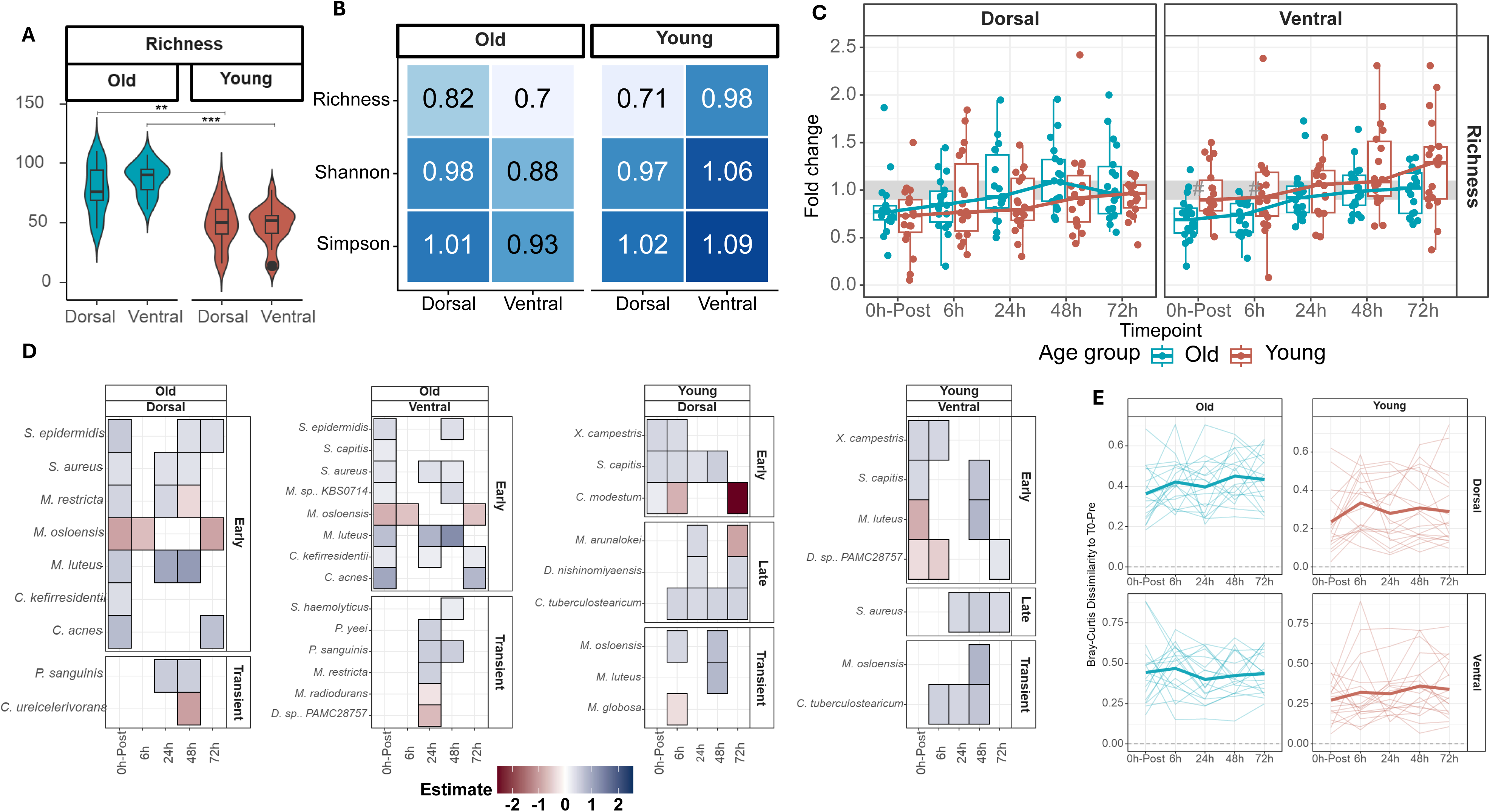
Skin microbiome recovery post tape-stripping. (A) Violin plots with inset boxplots showing the distribution of baseline richness values in older and younger age groups. Paired Wilcoxon tests were used for within-group comparisons, and Wilcoxon rank-sum tests for between-group comparisons. False discovery rate (FDR)–adjusted p-values are denoted as “***” for *p*≤0.001 and “**” for *p*≤0.01. In all boxplots, center lines denote medians, box limits the upper and lower quartiles, and whiskers the minimum and maximum values. (B) Heatmap showing fold-changes in indicated parameters between tape-stripped (TS) and non–tape-stripped (NTS) sites in younger and older age groups. (C) Boxplots showing the recovery of richness values across timepoints in both age groups on dorsal (left) and ventral (right) sites, respectively. In all boxplots, center lines denote medians, box limits the upper and lower quartiles, and whiskers the minimum and maximum values. (D) Heatmaps showing organisms that were differentially abundant following tape-stripping across timepoints for various sites and age groups. (E) Line plots depicting Bray–Curtis dissimilarity between microbiome profiles at each time point relative to the pre-stress baseline, stratified by age group and anatomical site. Individual trajectories are shown in thin lines, with the group median highlighted as a thicker line.

Despite consistencies in microbiome profiles, tape stripping induced distinct patterns of diversity loss in dorsal and ventral sites (t=0 pre vs post), with elderly subjects observing greater loss in ventral sites and younger subjects in dorsal sites (**Figure 3B**). Overall community compositions changed from pre-stripping to post-stripping (t=0) to an extent that was similar for dorsal and ventral sites in both age groups (**Supplementary Figure 8**), indicating that diversity losses were primarily in the form of loss/gain of low-abundance members (also seen in low Simpson diversity changes; **Figure 3B**). Correspondingly, these losses were made up for quite rapidly, with richness resembling pre-tape-stripping values frequently within 24 hours (fold-change>0.9; **Figure 3C**), and with evenness, Simpson, and Shannon indices exhibiting limited fold-change at all timepoints (>0.9, except for Shannon diversity in ventral sites for younger subjects and t<24h; **Supplementary Figure 9**).

Despite rapid recovery of diversity, the skin microbiomes of most individuals did not resemble the pre-tape-stripping community (**Supplementary Figure 10**). To identify if there were any consistent taxon-specific changes over the recovery time course, we applied a Linear Mixed-Effects (LME) modelling framework and grouped taxa based on their differential temporal patterns (**Methods**). While most taxa were specific to an age group or site, several were observed to be consistent between dorsal and ventral sites (e.g. *C. acnes*, *S. epidermidis*, *S. aureus*, *M. luteus* and *P. sanguinis* in old subjects, and *S. capitis*, *C. tuberculostearicum*, *M. luteus, X. campestris* and *M. osloensis* in younger subjects), and a few were shared across age groups (e.g. *M. luteus*, *M. osloensis*, *S. capitis* and *S. aureus*; **Figure 3D**). Many of the shared taxa represent early changes in the skin microbiome in abundant skin bacteria (e.g. *C. acnes*, *S. epidermidis*), likely representing robust shifts that track the recovery trajectory. No changes were specific to a site and shared across age groups, indicating that there were no strong site-specific signatures associated with recovery (**Figure 3D**). Instead, a few species such as *P. sanguinis* in older subjects and *M. osloensis* in younger subjects were consistently transiently enriched during the recovery process across sites, while frequent persistent enrichment of the pathobiont species *S. aureus* across sites and age groups could be a cause for concern in some individuals (**Figure 3D**).

To assess community-level recovery trajectories, we quantified Bray–Curtis dissimilarities between post-stress microbiomes and baseline microbiome configurations (**Method**). This analysis revealed substantial inter-individual heterogeneity, with wide variation in the trajectory and extent to which microbiomes return to their original states (**Figure 3E**). Recognizing that post-stress microbiomes may converge to alternative states^35^ rather than revert to baseline, we assessed temporal *stability* by computing Yue-Clayton theta index^36^ values between adjacent timepoints (**Methods**). Given the importance of microbiome stability for skin homeostasis^37^ and health^38^, our analysis revealed several notable trends: (i) >15% of sites did not achieve microbiome stability within our experimental timeframe (**Supplementary Figure 11A**), and (ii) when stability was attained, it was frequently site-specific (>35% of the subjects; **Supplementary Figure 11B**). Collectively, these findings highlight that while skin microbiomes rapidly recover diversity following skin stress, they do not often revert to their original composition or attain stability even after 3 days.

### Microbiome stability trajectories are associated with host attributes and microbial function

To investigate heterogeneity in microbiome stability trajectories, we clustered time-series profiles based on the Global Alignment Kernel distance (**Methods**) to reveal six distinct trajectory groups, each characterized by a unique temporal pattern of compositional similarity (**Figure 4A**). Notably, each cluster contained a mixture of trajectories from both younger and older subjects, and from dorsal and ventral skin sites, indicating that microbiome stability patterns are not solely determined by age or anatomical location (**Supplementary Figure 12**). Furthermore, these recovery patterns could be categorized into three broad types, each containing two distinct subgroups: (a) *Unperturbed*, characterized by minimal changes in stability across timepoints (U-H, U-L); (b) *Improved stability*, where stability increases relatively over time (IS-H, IS-L); and (c) *Reduced stability*, where stability is low in the final timepoints (RS-H, RS-L; **Figure 4A**).

**Figure 4.**
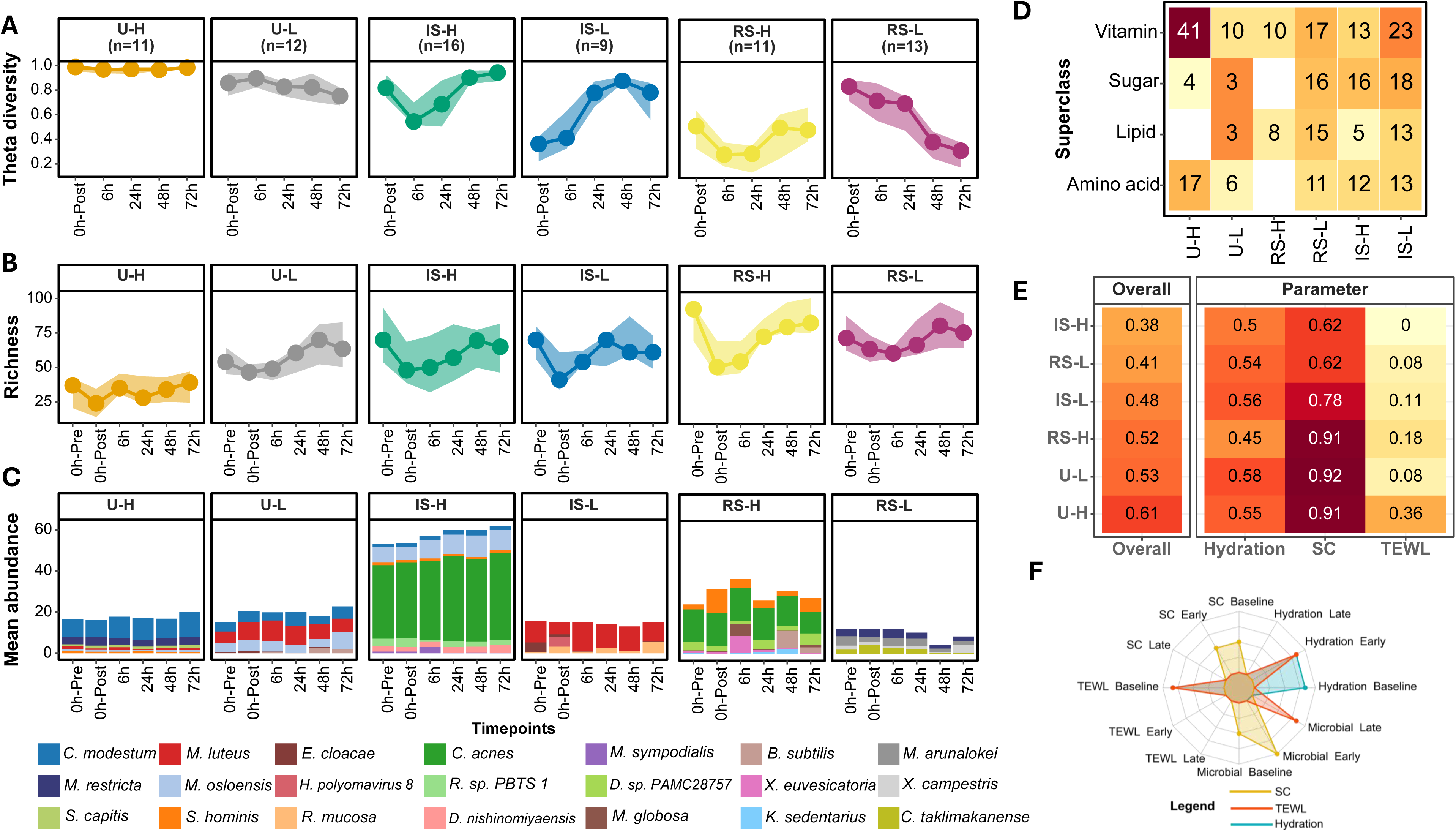
Trajectories of skin microbiome stability and associated functional features. (A) Line plots showing microbial stability trajectories across timepoints in each of the six defined groups (shaded areas depict 95% confidence interval). (B) Line plots of alpha diversity (richness) over time within the six stability groups (shaded areas depict 95% confidence interval). (C) Stacked bar plots depicting species-level composition across time points for each group; only the discriminating species are shown and are color-coded as shown in the legend. (D) Heatmap displaying the number of significantly enriched pathways within each stability group across all timepoints, grouped by functional superclass. (E) Heatmap showing the fraction of individuals who exhibited recovery for different parameters within each of the six stability groups. (F) Radar plot illustrating the relative influence of early (6–24h), late (48–72h), and baseline (pre- and post-stress) features on the prediction of 72h values for stratum corneum thickness (yellow), TEWL (red), and hydration (blue). Values represent normalized feature contributions within each recovery stage, visualized on a bounded proportional scale (0–0.4) to highlight relative dominance patterns across the three modalities.

Within the unperturbed trajectories, the U-H subgroup maintained consistently high stability values (median>0.9) across timepoints (**Figure 4A**) and was enriched for dorsal samples from younger subjects. Despite having the least richness, Shannon and Simpson diversity (**Figure 4B**, **Supplementary Figure 13**)—community composition was remarkably stable, defined only by minor variations in *C. modestum*, *M. restricta* and *S. capitis* (**Figure 4C**). Microbial functional profiling identified enrichment of several pathways involved in vitamin and cell wall biosynthesis, particularly L-lysine, L-histidine, L-arginine and L-threonine biosynthesis (across all timepoints) with the potential to support stratum corneum repair^39^ (**Figure 4D**, **Supplementary Figure 14, Supplementary Data File 5**). Intriguingly, this highly specialized community also associates with one of the highest rates of stratum corneum (SC) recovery (>90%; **Figure 4E**, **Supplementary Figure 15**). In contrast, the U-L subgroup is characterized by a flat but lower theta trajectory (median ∼0.8), and is enriched for samples from ventral sites and younger subjects (**Figure 4A**, **Supplementary Figure 12**). Here, richness was higher and largely unchanged after tape stripping, with taxonomic contributions from *M. luteus* and *M. osloensis* (**Figure 4B-C**). This group displayed slightly higher fraction of individuals recovering in terms of hydration measurements (>55%; **Figure 4D, Supplementary Figure 15**) with late-stage enrichment in L-tyrosine and L-phenylalanine microbial metabolism, highlighting its distinctness in host and microbial phenotypes relative to the U-H subgroup (**Supplementary Data File 5**).

Among the *Improved stability* subgroups, IS-L has a pattern of increasing stability after tape-induced stress in predominantly ventral samples from both age groups (**Figure 4A**, **Supplementary Figure 12)**. This group exhibits a functional enrichment in production of short-chain fatty acids (acetate, lactate, isobutanol) and an increase in *M. luteus* abundance, which together have been shown to play a role in promoting skin hydration^40, 41^ (**Figure 4C-D**, **Supplementary Figure 14, Supplementary Data File 5**). Coincidentally, this group also exhibits a higher rate of recovery for hydration measurements (>55%; **Figure 4E**). Finally, other subgroups such as IS-H, RS-H and RS-L are all marked by distinct recovery profiles for community richness, signature species, functional potential and skin physiological parameters (**Figure 4B-E**), highlighting the utility of this clustering for stratifying subjects and sites, and developing hypotheses for host-microbial interactions during post-stress recovery.

We next asked if the recovery of skin physiological parameters at the final timepoint (72h) could be predicted based on baseline and early timepoint features with machine-learning models (**Methods**). Our results suggest that TEWL recovery can be predicted the best (mean absolute error <5% for 0-1 scaled values) based on a combination of microbial features at later timepoints and baseline and early TEWL values (**Figure 4F**, **Supplementary Figure 16**, **Supplementary Data File 6**). In contrast, stratum corneum thickness and hydration were predicted with more modest accuracy, suggesting greater inter-individual variability. Notably, the inclusion of microbial features improved predictive performance across parameters, supporting a functional link between microbiome dynamics and host barrier recovery **(Figure 4F**, **Supplementary Figure 16**, **Supplementary Data File 6)**.

To further examine interdependencies between host physiology and microbial factors influencing recovery, we performed Cox proportional hazards analysis to model time to recovery across multiple parameters (**Methods**). Recovery timing following barrier disruption was jointly associated with early host physiological state, microbial community features, and stability trajectory membership, indicating that no single factor alone determined recovery dynamics (**Figure 5**). As expected, while physiological status in early timepoints had an impact on recovery profiles (e.g. higher values of TEWL predicts delayed TEWL recovery), microbial species abundances often had comparable influences, and in some cases stronger effects (e.g. *M. restricta* abundance in relation to TEWL recovery). Similarly, while TEWL values impacted hydration recovery in the model as expected, microbial taxa such as *S. hominis*, *M. arunalokei* and *C. modestum* also exhibited strong influences (**Figure 5**). In contrast, SC recovery appeared to have more selective influences including membership in distinct recovery trajectories (RS-L, IS-H) linked to delayed SC recovery, consistent with their observed association with lower SC thickness recovery rates (**Figure 4E**). Intriguingly, age group and body site also influenced recovery times in this model, despite not showing significant associations independently at the population level (**Figure 2A**), suggesting that their impact is more pronounced in combination with the microbial features modelled here. Finally, even after accounting for individual host and microbial taxa predictors, membership in various stability groups and community diversity values were associated with recovery of skin physiological parameters and the microbiome. Together, these analyses highlight that skin recovery following barrier disruption reflects combined effects of host physiology, microbial taxa/functions and microbiome recovery trajectories.

**Figure 5.**
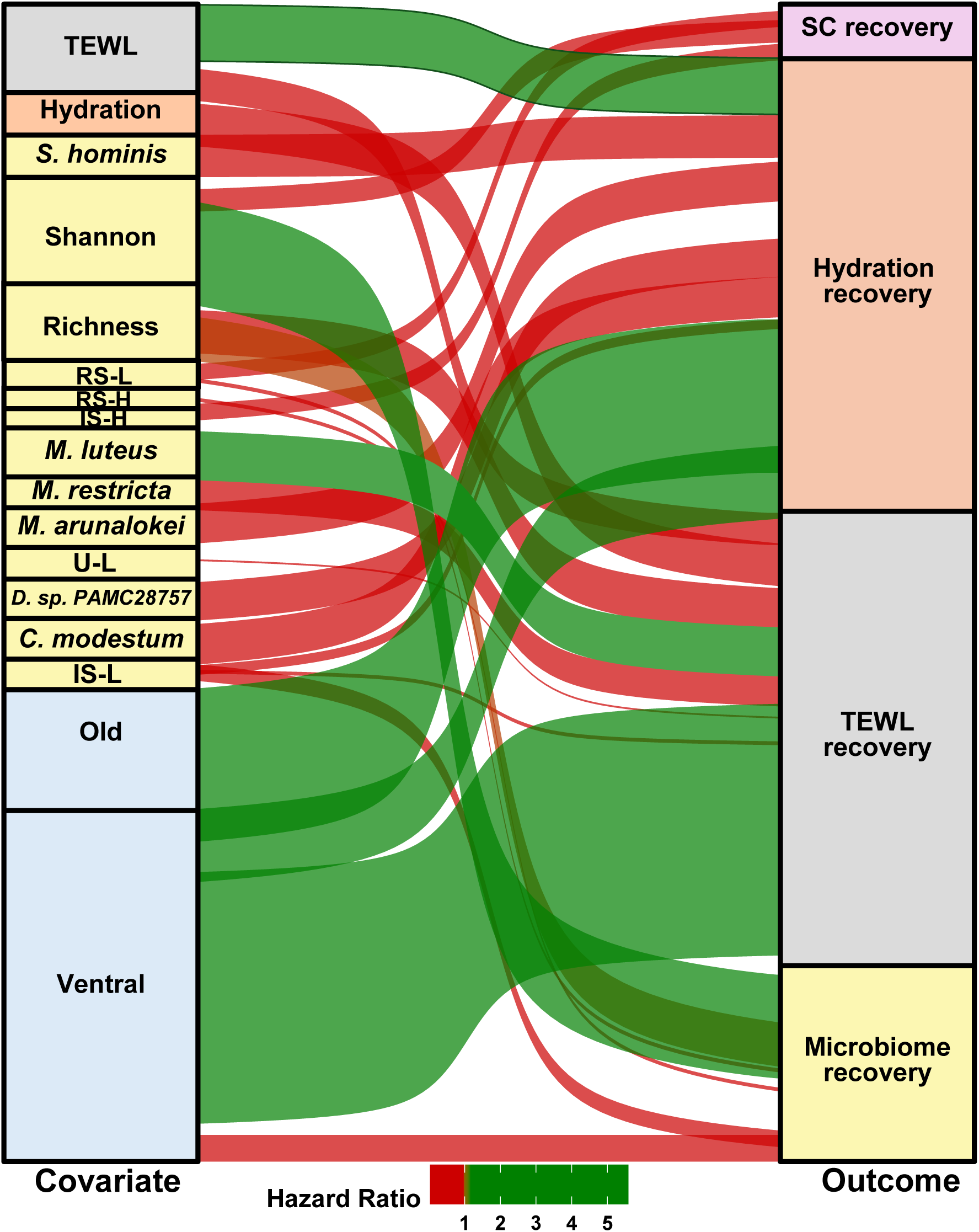
Multimodal predictors of recovery timing revealed by Cox proportional hazards modelling. Alluvial plot showing associations between early host physiology, microbial features, and stability trajectories with recovery of TEWL, stratum corneum thickness, hydration, and microbiome composition. Flow width reflects the relative strength of association, while colour indicates directionality, with red denoting reduced hazard (delayed recovery) and green denoting increased hazard (accelerated recovery). For hazard ratios <1, narrower flows indicate stronger delaying effects, whereas for hazard ratios >1, wider flows indicate stronger effects. All associations are statistically significant with FDR-adjusted *p*<0.05.

## Discussion

Human skin is continuously exposed to a wide range of environmental, mechanical, and biological stressors, yet must rapidly restore barrier integrity to maintain homeostasis and prevent infection. Although skin aging is commonly associated with impaired repair capacity^42, 43^, resilience is increasingly recognized as a context-dependent property shaped by anatomical location, baseline physiology, and interactions with resident microbial communities^44–46^. Understanding how these factors affect recovery following acute stress remains a fundamental knowledge gap in skin biology^47^.

In this study, we investigated how age, anatomical site, and the skin microbiome are associated with recovery patterns following acute mechanical barrier disruption. By combining longitudinal measurements of skin structure, physiology, and microbial composition with trajectory-based analyses of microbiome stability, we show that recovery from a standardized insult is highly heterogeneous across individuals. Through multimodal data integration, we uncovered that recovery trajectories diverge early and are shaped by specific combinations of host physiological and microbiome features, rather than individual variables considered in isolation. Across both age groups and anatomical sites, we observed a broadly consistent recovery trajectory, wherein a majority of individuals achieved a near-complete restoration of SC structure, yet exhibited persistently impaired TEWL recovery. This apparent decoupling between structural and functional restoration likely reflects incomplete reconstitution of lipid organization and intercellular junction integrity^8, 48^, processes that may not be fully captured by gross structural measures such as SC thickness. Indeed, the TEWL recovery trajectory displayed a distinct pattern, with consistently higher fold changes observed in younger individuals relative to older counterparts, particularly on the dorsal side. Such site- and age-dependent variability in TEWL likely reflects differences in baseline barrier function and susceptibility to perturbation^49, 50^. Furthermore, frequent co-recovery of SC thickness and hydration, in contrast to TEWL, supports the notion that these parameters reflect coordinated structural reconstitution of the epidermis^51, 52^, whereas TEWL captures the functional aspects of barrier integrity^53^.

Acute mechanical disruption of the skin barrier also induced immediate and structured perturbations to the skin microbiome, consistent with prior studies showing that tape stripping removes the outermost stratum corneum together with surface-associated microbial taxa^26, 28, 54^. We observed immediate reductions in richness and diversity, characterised by transient loss of low-abundance taxa and selective expansion of stress-tolerant or opportunistic organisms, consistent with perturbation-driven restructuring reported across other host-associated microbial ecosystems^55, 56^. Notably, several taxa exhibited sustained, site- and age-dependent responses, indicating that microbial reassembly is context dependent. For example, consistent enrichment of *S. capitis* following stress suggests a role for competitive or antimicrobial traits in rapid recolonization^57^, whereas depletion of resident taxa such as *C. modestum,* particularly on young dorsal skin may reduce functional capacity during early recovery^58^. Notably, we observed persistent increases in *S. aureus* in older individuals and *C. tuberculostearicum* on the young dorsal skin. Expansion of such taxa, often associated with barrier dysfunction or inflammatory skin conditions, may indicate a shift toward stress-adapted or dysbiosis-prone configurations^59, 60^. Furthermore, our longitudinal data revealed pronounced heterogeneity in microbial recovery following perturbation: while some individuals’ microbiomes returned close to their pre-stress configuration, others stabilized in alternative compositional states that diverged from baseline, supporting the presence of multiple recovery trajectories following barrier disruption (**Figure 3E**).

Such heterogeneous outcomes are a hallmark of perturbed ecological systems and indicate that recovery cannot be adequately described by simple return-to-baseline metrics^35, 56, 61^. Instead, these observations motivated a stability-based clustering approach to capture whether distinct microbial configurations, and their temporal trajectories, differentially support recovery of host physiological parameters. By clustering individual time-series profiles of microbial stability, we identified reproducible recovery trajectories that reflected distinct modes of post-stress reorganization rather than uniform restoration. Importantly, these trajectories were shared across age groups and anatomical sites but differed in their relative enrichment, highlighting that host context biases the likelihood of entering specific recovery states.

This trajectory-based clustering revealed that microbial stability is not inherently beneficial or detrimental. Unperturbed trajectories, particularly U-H, which was enriched for young dorsal samples, were characterized by low diversity but strong dominance of a restricted set of taxa, including *C. modestum*, *Malassezia restricta*, and *S. capitis*, alongside enrichment of amino acid and vitamin biosynthesis pathways. These communities coincided with robust stratum corneum recovery, suggesting that functionally aligned^39^, low-diversity states can still help in building resilience. In contrast, other trajectories exhibiting stable or rebounding microbial configurations were associated with incomplete physiological recovery, despite apparent compositional stabilization. This highlights that stabilization into a non-restorative microbial state, rather than instability *per se*, can underlie poor recovery. Taxa associated with lipid metabolism, including shifts in *Malassezia* species composition (*M. restricta*, *M. arunalokei*), were particularly prominent in trajectories linked to impaired hydration recovery. Given the central role of *Malassezia* in sebum utilization^62^ and lipid processing^63^, altered fungal community structure may disrupt lipid-mediated barrier repair and moisture retention, either through functional involvement or as a marker of underlying barrier state. Conversely, trajectories enriched in short-chain fatty acid–producing pathways and taxa such as *M. luteus* were associated with improved hydration recovery, consistent with emerging evidence linking microbial metabolites to skin hydration and barrier function^40, 64, 65^.

Collectively, these organism-level patterns support a model in which recovery failure arises not from a lack of microbial stability, but from stabilization into microbial configurations that may not be well aligned with host repair processes. Rather than a uniform trajectory, individuals exhibit different microbiome states, with some stabilizing into resilient, functionally supportive configurations (such as IS-L), while others transition into low stability states (such as RS-L) marked by compositional shifts in key skin commensals, particularly differential abundance of *Malassezia* species, coupled with reduced stratum corneum restoration. Consistent with this heterogeneity, age and anatomical site were associated with specific microbiome stability trajectories, but recovery outcomes diverged markedly among individuals within the same age-site category.

Finally, our predictive modelling demonstrates that early host and microbial features can prospectively inform recovery outcomes, with TEWL emerging as the most predictable parameter and microbial features contributing to stratum corneum and hydration recovery. Cox survival analysis further supported this framework by showing that the timing of recovery is shaped by early host physiology and microbial state rather than by age or magnitude of stress alone. These findings suggest that early post-stress microbial composition**—**and the specific taxa that dominate during this window—may serve as actionable indicators or intervention targets to redirect recovery toward desired trajectories. Future studies involving targeted interventions, such as microbiome-based therapeutics, can help evaluate whether shifts in microbial configurations toward functionally supportive states enhance barrier resilience. Complementary *in vitro* experiments using reconstructed skin models or keratinocyte cultures, can further delineate the causal roles of specific taxa.

Together, our findings show that recovery from skin barrier disruption is a heterogeneous, trajectory-dependent process shaped by early host physiology and microbial reorganization rather than age alone. By integrating microbiome changes with host physiology, this work highlights the importance of early host–microbiome relationships in determining recovery outcomes and identifies microbial configurations as potential targets for enhancing skin resilience following stress.

## Methods

### Study design

The study was approved by National Healthcare Group Domain Specific Review Board (2019/01127, 2022/00628), Singapore. Subjects provided additional data via a questionnaire, with questions covering general skin and skin health. Exclusion criteria for subjects were any self-reported skin condition and current antibiotic usage. The pilot study included 10 healthy Singaporean Chinese female subjects (mean age 40 ± 3 years; Fitzpatrick phototype II–III). In the current study, a total of 36 healthy Singaporean Chinese female subjects (Fitzpatrick Photo-type II-III) were enrolled; 18 subjects were aged 21-31 (‘young’ group) and another 18 subjects were aged 55-65 (‘old’ group). A schematic showing the design for the study is provided in **Figure 1A**.

Subjects were provided with “le soin lavant toleriane” and were advised to use the body wash, starting two days prior to the study start date until the end of the study. Each subject was acclimatised to the clinical environment at least for 30 minutes prior to sample collection, and data collection timepoints were kept relatively uniform. On the first day of sampling, a temporary adhesive mark was applied to guide sample collection and image acquisition. Sites were randomised for tape-stripping and non-tape-stripped adjacent controls (**Figure 1A**). The skin stress procedure was implemented via tape stripping with a 22mm diameter D-Squame tape (Clinical & Derm) with controlled pressure applied using D500-Dsquame pressure instrument till a 2× increase in TEWL was achieved. A new D-Squame tape was used after each tape stripping action.

Eight sample sites in sun exposed (dorsal) and non-exposed (ventral) forearm, were used in this study (**Figure 1A**). Sampling locations were delineated as either non-tape stripped sites (NTS), which served as controls, or as tape stripped sites (TS). Physiological and imaging measurements were obtained from the TS and NTS sites at each time point. Among the NTS sites, one site was randomly selected at each time point for physiological measurements, while the other was designated for imaging. For microbiome profiling, tape-strip samples were collected from one randomly selected site among five pre-marked locations at each time point. A total of 432 skin D-Squame tapes (green region), were collected from 36 subjects these samples were collected from control and pre-stripped target skin sites at six different time points (pre- and post-tape stripping on day 0 as well as at 6-, 24-, 48- and 72-hours post-stress; **Figure 1A**). Tapes were placed in a microcentrifuge tube and stored at −80°C. Separately, two specific sites were set aside for the ms-RSOM image acquisition to ensure no impact on microbiome sample collection (region showed in pink in **Figure 1A**).

### Skin physiology measurements

Three skin physiology measurements were made, including TEWL (VapoMeter), skin hydration (MoistureMeterSC) and elasticity (ElastiMeter) [all Delfin devices]. The devices were acclimatized and calibrated before use. Skin physiological parameters were measured at all time points during the study. Each device was applied to the skin until the measurement was taken, with three measurements being taken at each time point and averaged for further analysis.

### Skin structural measurements using multispectral optoacoustic mesoscopic imaging (ms-RSOM)

The ms-RSOM system (RSOM Explorer ms-C50, iThera Medical GmbH, Munich, Germany) integrates a nanosecond Raman laser with four distinct wavelengths (532, 555, 579, and 606 nm) and an ultra-broadband transducer (centre frequency 50 MHz, bandwidth 11-99 MHz). The absorption of light by different chromophores leads to the production of pressure waves, detectable through an ultrasound transducer, subsequently reconstructed into single-wavelength images. To enhance visualization, a frequency separation approach was implemented such that the lower frequency band (spanning 11 to 33 MHz) was used to study to larger structures, while the higher frequency band (ranging from 33 to 99 MHz) enabled detection of smaller vascular structures^66^. Spectral unmixing was performed using a linear regression algorithm incorporating non-negative constraints to discern distinct skin chromophores based on their respective absorption spectra within the induced light wavelengths. Illumination at multiple wavelengths generated a three-dimensional spatial map of skin chromophores, including melanin, HbO_2_ and Hb through spectral unmixing. This also allowed for the quantification of functional information, such as sO_2_ in microvasculature. Specific metrics derived from the ms-RSOM images included (1) stratum corneum (SC) thickness, (2) epidermis thickness, (3) melanin thickness, (4) sO_2_, and (5) total blood volume (TBV). Quantitative analysis of each metric was conducted for all subjects before and after tape stripping.

### Calculation of imaging-based metrics

We devised an in-house automatic segmentation algorithm to compute skin metrics. Utilizing the 532nm excited optoacoustic profile along the skin depth, we delineated the boundary between the epidermis and dermis. This boundary was determined by averaging all pixels in the x-y plane along the depth (z-axis) from the 3D image stacks. The skin surface was identified as the starting point of the high frequency band of the optoacoustic profile along the depth.

To ascertain the centre of the melanin layer, which strongly absorbs 532nm light, we identified the overlap of the first dominant peak between the low and high frequency bands of the optoacoustic profile. Using the full width at half maximum (FWHM), we determined the edges of the melanin layer, with the distance between these edges considered as the thickness of the melanin layer. The distance from the skin surface to the upper edge of the melanin layer was designated as SC thickness, as the SC comprises the outer layer of the epidermis. Conversely, the distance from the skin surface to the lower edge of the melanin layer was regarded as the epidermis thickness in our study, as melanin resides in the basal layer of the epidermis.

sO_2_ levels were determined by the equation HbO_2_/(Hb+HbO_2_), and the contributions of HbO_2_ and Hb in the dermis were obtained through spectral unmixing based on the absorption coefficients of skin chromophores from individual wavelength images. TBV was derived by summing the non-zero number of voxels within the segmented 3D dermal region after applying a threshold.

### Definition of recovery for skin physiological and imaging parameters

Recovery was quantified using fold-change values relative to the baseline for each subject and anatomical site obtained at the initial timepoint. An observation was considered recovered if the fold-change value fell within a predefined range of 0.9–1.1, indicating a return to baseline levels. Group-level recovery was determined using statistical comparisons, where non-significant differences from baseline (FDR-adjusted *p*≥0.05) indicated recovery at the cohort level. Recovery in individual subjects were classified into three categories: Recovered (recovery observed at the final timepoint or sustained over at least two consecutive timepoints), Partially recovered (recovery observed at one or more isolated timepoints), and Non-recovered (no recovery observed across all timepoints). Recovery time was defined as the earliest timepoint at which recovery occurred.

### Genomic DNA extraction

Genomic DNA was extracted from skin tapes via a bead beating step on a FastPrep-24 Automated Homogeniser (MP Biomedicals), followed by magnetic bead based extraction using an EZ1 Advanced XL Instrument (Qiagen) with EZ1 DNA Tissue Kit (Qiagen). Specifically, 500µl of Buffer ATL (Qiagen) was added to each sample transferred into a Lysing Matrix E tube. Sample tubes were bead-beated at4 m/s for 30 seconds, twice. After homogenization, tubes were centrifuged at maximum speed for 5 minutes, with 200µl of the resulting supernatant transferred into 2ml EZ1 sample tubes (Qiagen). 12µl of Proteinase K was added to the supernatant, followed by vortexing and incubation at 56°C for 15 minutes. Finally, samples were transferred to the EZ1 Advanced XL Instrument (Qiagen) for purification, with a final eluate of 50µl in buffer EB. Extracted DNA samples were stored in 1.5ml elution tubes at −20 °C. As a negative control, a blank tape was transferred into a Lysing Matrix E tube and processed in the same way.

### Construction and sequencing of metagenomic libraries

26µl of purified genomic DNA underwent NGS library construction steps using NEBNext Ultra II FS DNA Library Prep Kit according to manufacturer’s instructions. Purified DNA was added to fragmentation reagents, mixed well and placed in a thermocycler for 10 mins at 37°C, followed by 30 mins at 65 °C to complete enzymatic fragmentation. Post fragmentation, adaptor-ligation was performed using Illumina-compatible adaptors, diluted 10-fold as per kit’s recommendations prior to use. Post-ligation, size selection or cleanup of adaptor-ligated DNA was performed using Ampure XP beads in a 7:10 beads-to-sample volume. Unique barcode indexes were added to each sample and amplified for 12 cycles under recommended kit conditions to achieve multiplexing within a batch of samples. Samples were purified using Ampure XP beads in a 7:10 beads-to-sample volume, with a final elution volume of 20µl. Finally, each library sample was assessed for quality based on fragment size and concentration using the Agilent D1000 ScreenTape system, and quality-checked samples were adjusted to identical concentrations by means of dilution and volume-adjusted pooling. For sequencing, 74-78 samples (including one negative control) were pooled together for one pooled library. The multiplexed sample pool was paired-end (2×151bp) sequenced on the Illumina HiSeq X Ten platform to generate an average of 9 million reads per sample. All sequencing was done in the Novogene sequencing facility in Singapore in accordance with standard Illumina sequencing protocols.

### Read pre-processing and taxonomic profiling

Shotgun metagenomic sequencing reads were processed using a Nextflow pipeline (https://github.com/CSB5/shotgunmetagenomics-nf). Briefly, raw reads were filtered to remove low quality bases and adapter sequences were removed using fastp^67^ (v0.20.0) with default parameters. Human host reads were removed by mapping to the hg38 reference using BWA-MEM^68^ (v0.7.17-r1188, default parameters) and samtools^69^ (v1.9) with parameters -f12 -F256. The remaining reads were profiled using Kraken2^70^ (v.2.0.8) and Bracken^71^ (v.2.6.1) for taxonomic abundances. The database^72^ used was a 50Gb Kraken2 database built from Refseq bacterial, archaeal, viral and fungal genomes, plasmid sequences and the hg38 human reference genome. This database also contains additional Malassezia assemblies downloaded from NCBI.

### Kitome identification

Potential reagent-derived contaminants (kitome) were identified using a multi-criteria approach integrating negative controls, library DNA concentration, and compositional correlation analysis. Taxa present in negative control libraries were first used to define candidate contaminants. Associations between taxa abundance and library concentration were assessed using Spearman correlation, with taxa showing significant negative correlations indicative of low-biomass contamination. To further refine contaminant identification, we performed correlation analysis using CCREPE (v1.36.0) and classified taxa exhibiting strong correlations with blank-associated taxa (ρ > 0.65, adjusted p-value < 0.1) as potential contaminants. The final kitome set included taxa enriched in blanks, negatively associated with library concentration, or strongly correlated with blank-associated taxa. To account for shared contamination sources at higher taxonomic levels, additional taxa belonging to the same genera as identified contaminants were also included. Identified kitome taxa were removed, and profiles were renormalized using total sum scaling prior to downstream analyses.

### Diversity and differential abundance analysis

Species level taxonomic profiles were used to compute alpha diversity indices including richness, Shannon and Simpson diversity and a beta diversity index (Bray-Curtis dissimilarity) using the R package vegan (v2.6.4). Taxonomic profiles were filtered by retaining only those taxa present at a mean abundance of more than 0.1% in at least 50 samples per site and per age group. Differential abundance analysis was carried out to identify timepoint based variations per group and per site using linear mixed effects modelling (lme) with random effects as subject IDs and log transformed filtered taxonomic profiles (nlme v3.1-168 package in R). Model fit was computed through estimated marginal means using the emmeans package in R and the FDR method was used to account for multiple testing. Taxa were classified into three temporal categories based on the time points at which they showed significant differential abundance: (i) “Early” taxa were significantly changed immediately after tape-stripping, (ii) “Late” taxa were significantly changed only at later timepoints and particularly at the last timepoint, and (iii) “Transient” taxa were all others that were only significantly different in intermediate timepoints.

### Definition of temporal microbiome stability

Microbiome stability was assessed using temporal dynamics of community composition based on Yue–Clayton (YC) similarity. For each subject and anatomical site, pairwise changes in YC similarity between consecutive timepoints were computed, and a stability threshold was defined as an absolute change ≤0.10. Sustained stability was identified by the presence of at least two consecutive stable intervals, minimizing the impact of transient fluctuations. To account for gradual re-establishment, an endpoint criterion was additionally applied, whereby samples with YC similarity ≥0.75 at later timepoints (≥24h) were considered stable. The earliest timepoint satisfying either sustained stability or endpoint criteria was defined as the time to microbiome stabilization and used for downstream cox survival analyses.

### Time-series clustering of microbiome trajectories

Unfiltered taxonomic profiles were used to compute the Yue–Clayton theta index between adjacent timepoints (stability values) using custom R scripts. The resulting stability values were organized as a time-series, and pairwise distances between trajectories were calculated using the Global Alignment Kernel (GAK)^73^ method. The GAK distance matrix was next subjected to partitional clustering, with cluster centroids defined by the median trajectory shape. The optimal number of clusters was determined using the means of normalised Silhouette score, Dunn Index and Davies-Bouldin Index. Cluster stability was assessed using adjusted rand index (ARI) and Jaccard index after bootstrapping 500 times.

### Taxonomic variation across stability clusters

To identify taxonomic features distinguishing various stability clusters, we first selected the ten most abundant features in each group (based on mean abundance). For these features, variance across timepoints was calculated and used as input for one-vs-rest classification. Discriminatory power was assessed by the Area Under the Receiver Operating Characteristic Curve (AUC), and taxa with AUC>0.79 were considered key discriminants.

### Pathway analysis

The HMP Unified Metabolic Analysis Network (HUMAnN3)^74^ pipeline was applied to quantify the relative abundance of microbial pathways in skin metagenomes, using filtered taxonomic profiles generated by Kraken2. Unstratified pathway abundance values across time points were then analysed to detect significant temporal changes between microbiome stability groups, with Linear Discriminant Analysis (LDA > 3) used to identify discriminative pathways.

### Machine learning framework for endpoint value prediction

To predict values at the final time point, we applied linear regression, random forest, and gradient boosting under two complementary strategies (**Supplementary Figure 17**). In the temporal modelling strategy, models were trained on all non-empty combinations of four feature categories (microbial, stratum corneum, TEWL, and hydration; 15 combinations), using either snapshot prediction with features from a single time point or cumulative prediction incorporating features from the current and previous time points. In the feature optimization strategy, random forest was used to rank features and identify the top ten across all time points, followed by best subset selection generating all possible subsets of size 1–10 (1,023 combinations). Model performance was evaluated using mean absolute error (MAE), and the best models were chosen based on lowest MAE, with preference given to models requiring fewer features. The contributing features were grouped into three temporal categories— Baseline (T0-Pre, T0-Post), Early (T1, T2), and Late (T3, T4)—to visualize their relative influence on prediction.

### Cox survival analyses

To quantify the contribution of individual covariates to parameter-specific recovery, we applied Cox proportional hazards modelling incorporating continuous features from four domains—microbial, transepidermal water loss (TEWL), stratum corneum (SC) thickness, and hydration—as well as categorical factors including age, body site, and microbial stability group. Microbial features comprised of alpha diversity metrics (Shannon, Simpson, and richness) and the five most abundant baseline species, stratified by age group and body site. Model performance was evaluated using the concordance index (C-index), and models with C-index values >0.75 were retained for downstream analyses. Separate Cox models were constructed for TEWL, SC thickness, and hydration recovery, with time-varying covariates included to account for longitudinal changes.

## Data Availability

Shotgun metagenomic sequencing data from this study is available from the European Nucleotide Archive (ENA – https://www.ebi.ac.uk/ena/browser/home) under project accession number PRJEB108618.

## Notes

### Competing Interest Statement

The authors have declared no competing interest.

### Funding

- A*STAR Research Entities: H22J1a0040
- LVMH (France)
- National Research Foundation: NRFI09-0015

