## Supplementary Figures for "Host recovery after skin barrier disruption is individual-specific and associated with microbial functions"

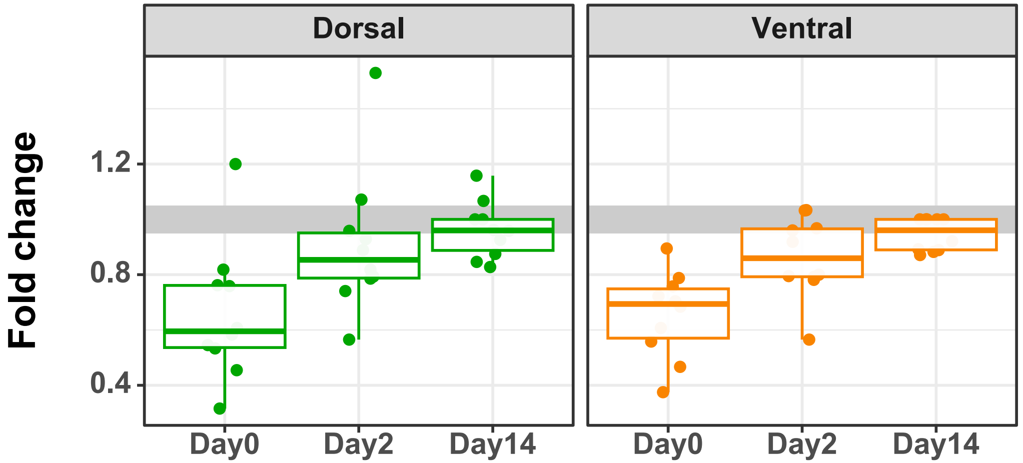


**Supplementary Figure 1 – Recovery of the stratum corneum following tape stripping.** Boxplots displaying the fold-change in stratum corneum thickness across dorsal and ventral sites at Days 0, 2, and 14 in the pilot study. Fold change values represent the ratio of tape-stripped (TS) to non–tape-stripped (NTS) measurements at each timepoint. The center line indicates the median, the box boundaries represent the interquartile range (IQR), and whiskers extend to the minimum and maximum values. Shaded gray box indicates the recovery region.

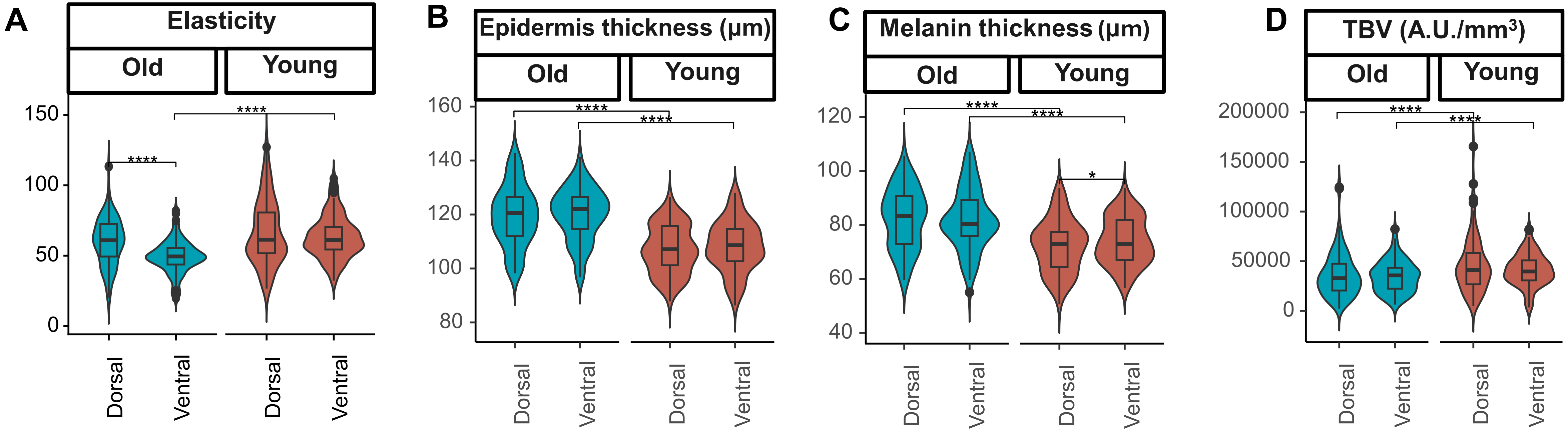


**Supplementary Figure 2 – Baseline differences between age groups and across sites:** Violin plots with inset boxplots showing the distribution of (A) Elasticity, (B) Epidermis thickness (μm), (C) Melanin thickness (μm), and (D) Total Blood Volume (A.U./mm$^3$) values on dorsal (left) and ventral (right) sites across all timepoints from the non–tape-stripped (NTS) side. Paired Wilcoxon tests were used for within-group comparisons, and Wilcoxon rank-sum tests for between-group comparisons. False discovery rate (FDR)–adjusted p-values are denoted as "****" for p≤0.0001 and "*" for p≤0.05. In all boxplots, the center line denotes the median, box limits the upper and lower quartiles, and whiskers the minimum and maximum values.

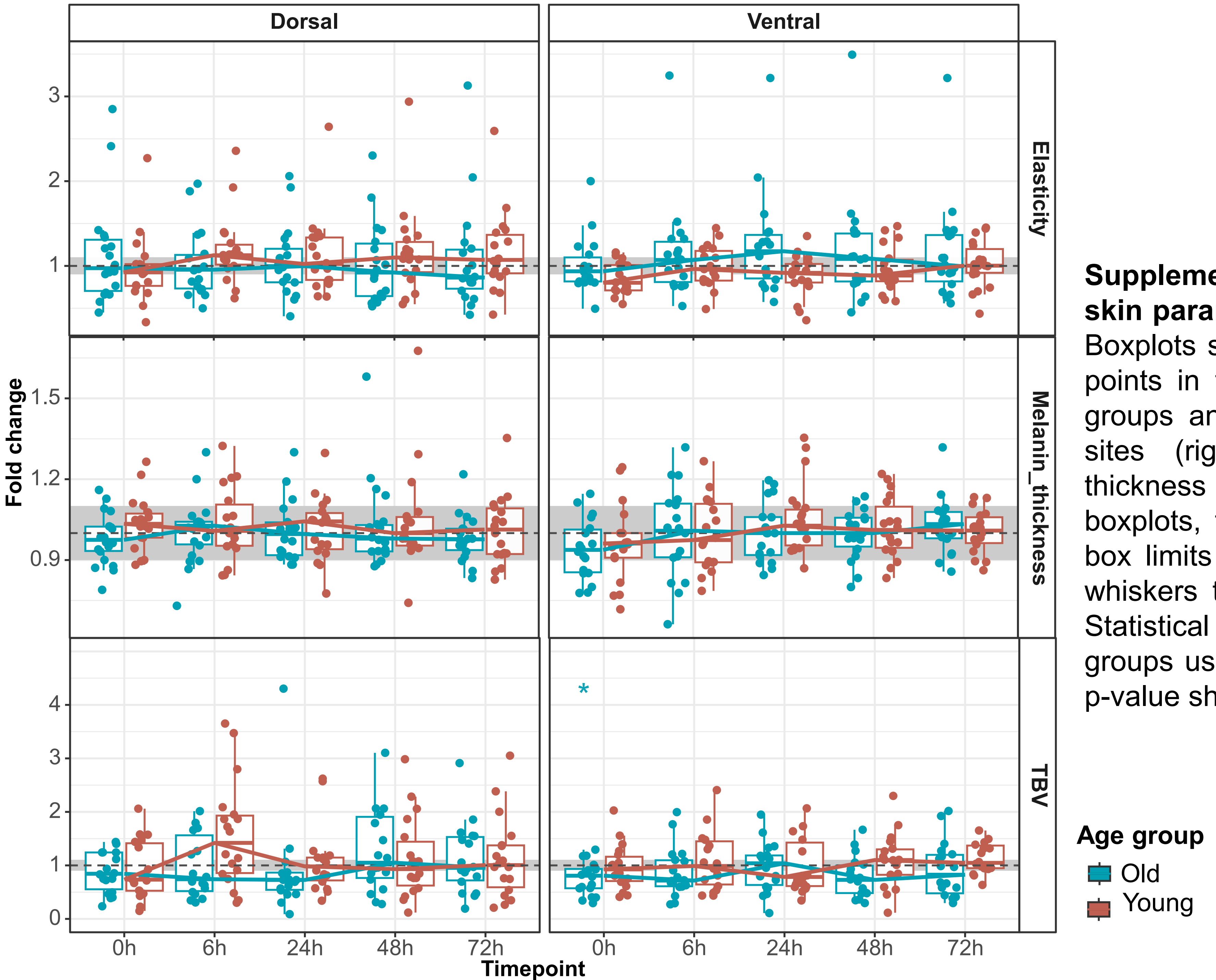


**Supplementary Figure 3 – Change in other skin parameters over time after skin stress.** Boxplots showing fold changes across all time points in the old (blue) and young (red) age groups and in dorsal (left panel) and ventral sites (right panel) for elasticity, melanin thickness and total blood volume (TBV). In all boxplots, the center line denotes the median, box limits the upper and lower quartiles, and whiskers the minimum and maximum values. Statistical tests were performed between age groups using Wilcoxon test with FDR adjusted p-value shown as "*" for p-value<0.05.

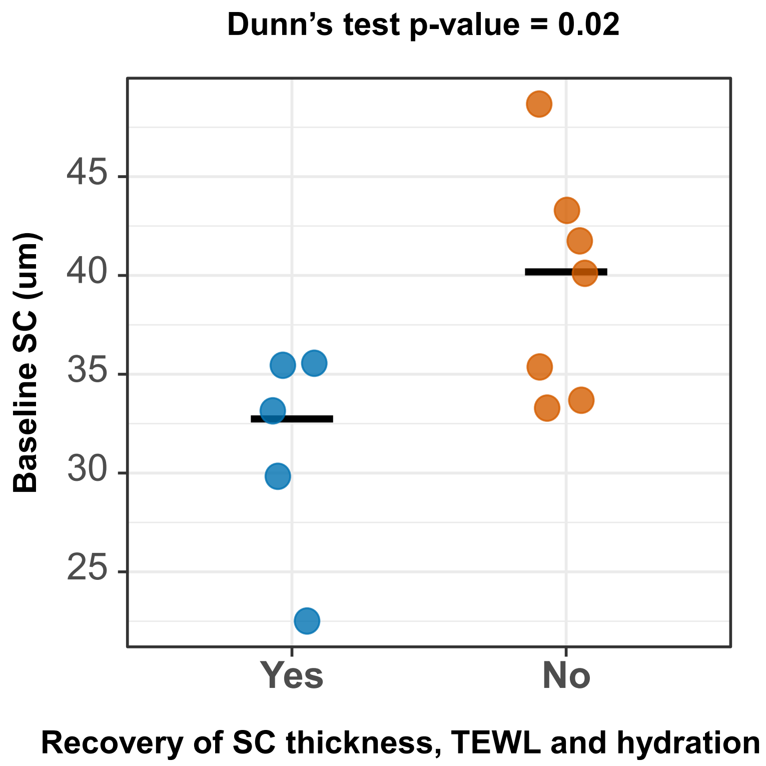


**Supplementary Figure 4 - Baseline stratum corneum differences between subjects who recovered across all parameters versus those who did not recover in any.** Plot showing baseline stratum corneum (SC) thickness measurements for subjects classified as recovered versus non-recovered based on all the TEWL, hydration, and SC outcomes. Points represent individual subjects, and crossbars indicate the median. Subjects who achieved recovery across all three physiological parameters exhibited lower baseline SC thickness values versus those who did not exhibit recovery for any of the parameters. Statistical differences were assessed using a Dunn's test, with the corresponding p-value shown.

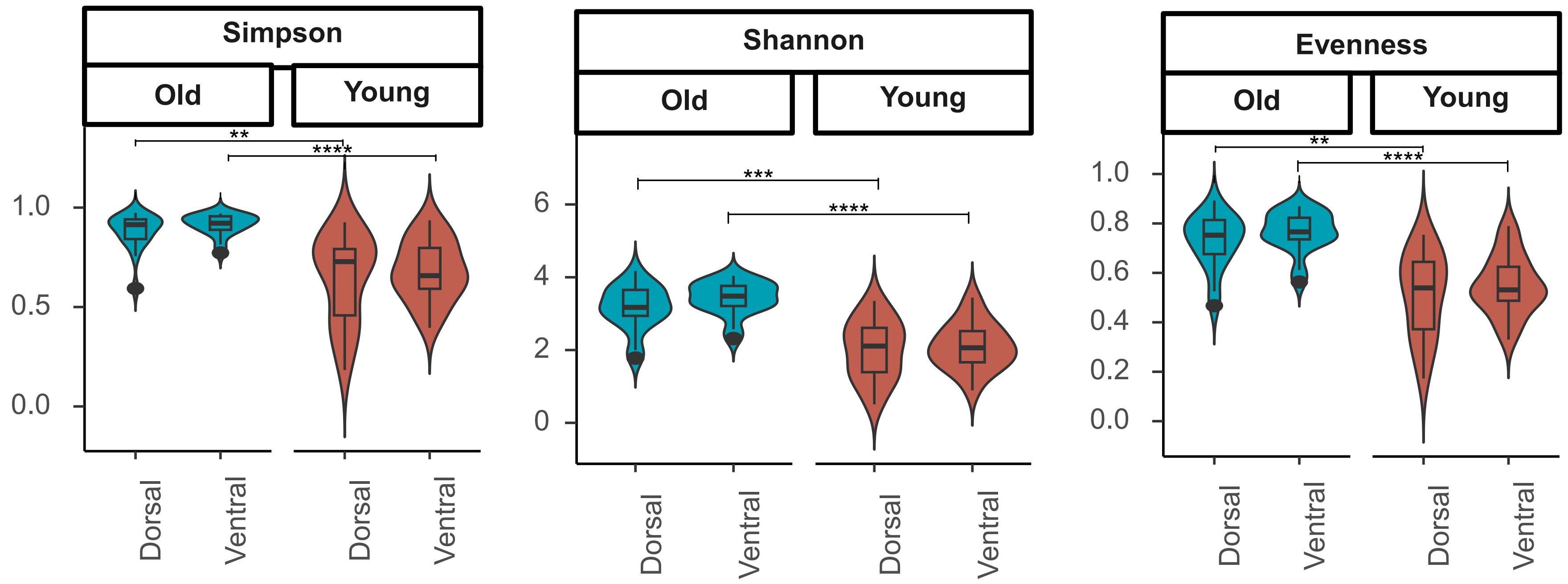


**Supplementary Figure 5 – Baseline differences in skin microbiome alpha diversities between the two age groups and across sites:** Violin plots with inset boxplots showing the distribution of Simpson diversity, Shannon diversity and Evenness values for dorsal (left) and ventral (right) sites across age groups from the non–tape-stripped (NTS) site. Paired Wilcoxon tests were used for within-group comparisons, and Wilcoxon rank-sum tests for between-group comparisons. False discovery rate (FDR)–adjusted p-values are denoted as "*****" for p≤0.0001, "***" for p≤0.001 and "**" for p≤0.01. In all boxplots, the center line denotes the median, box limits the upper and lower quartiles, and whiskers the minimum and maximum values.

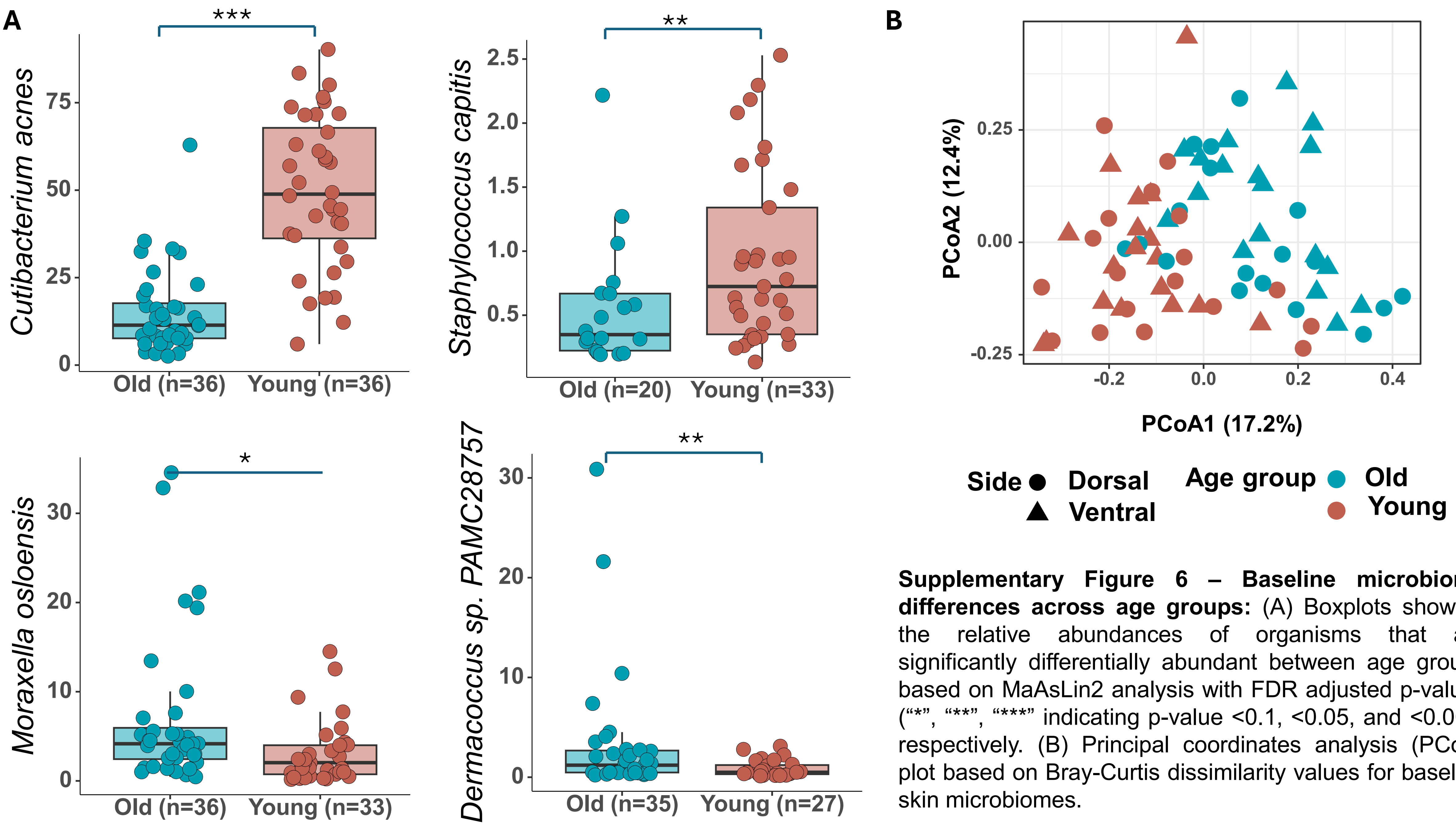


**Supplementary Figure 6 – Baseline microbiome differences across age groups:** (A) Boxplots showing the relative abundances of organisms that are significantly differentially abundant between age groups based on MaAsLin2 analysis with FDR adjusted p-values ("*", "**", "***" indicating p-value <0.1, <0.05, and <0.001, respectively. (B) Principal coordinates analysis (PCoA) plot based on Bray-Curtis dissimilarity values for baseline skin microbiomes.

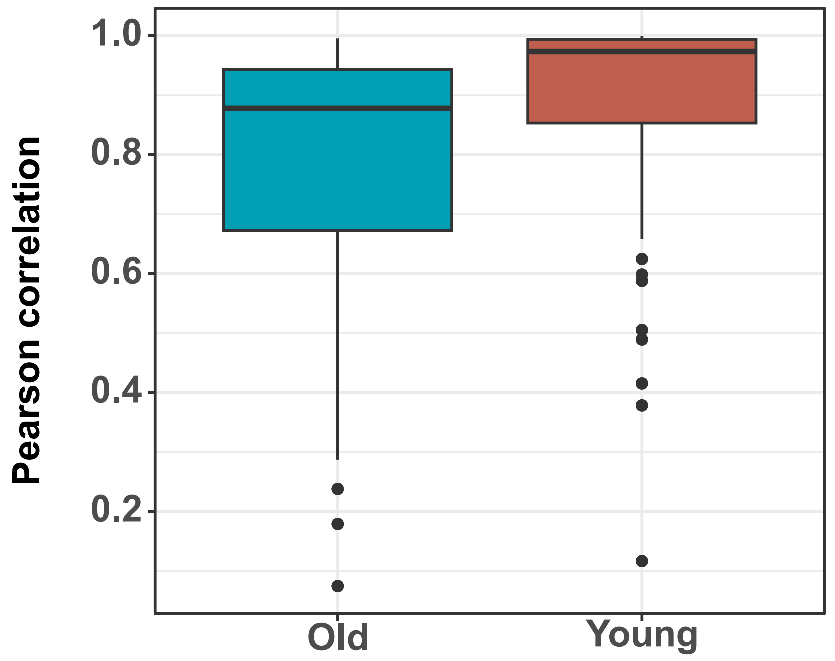


**Supplementary Figure 7 - Pearson correlation values between dorsal and ventral skin microbiomes.** Boxplots showing the distribution of Pearson correlation coefficients for the old and young age groups, indicating the degree of similarity in microbial community composition between dorsal and ventral sites. The center line denotes the median, box boundaries represent the interquartile range, and whiskers extend to the minimum and maximum values.

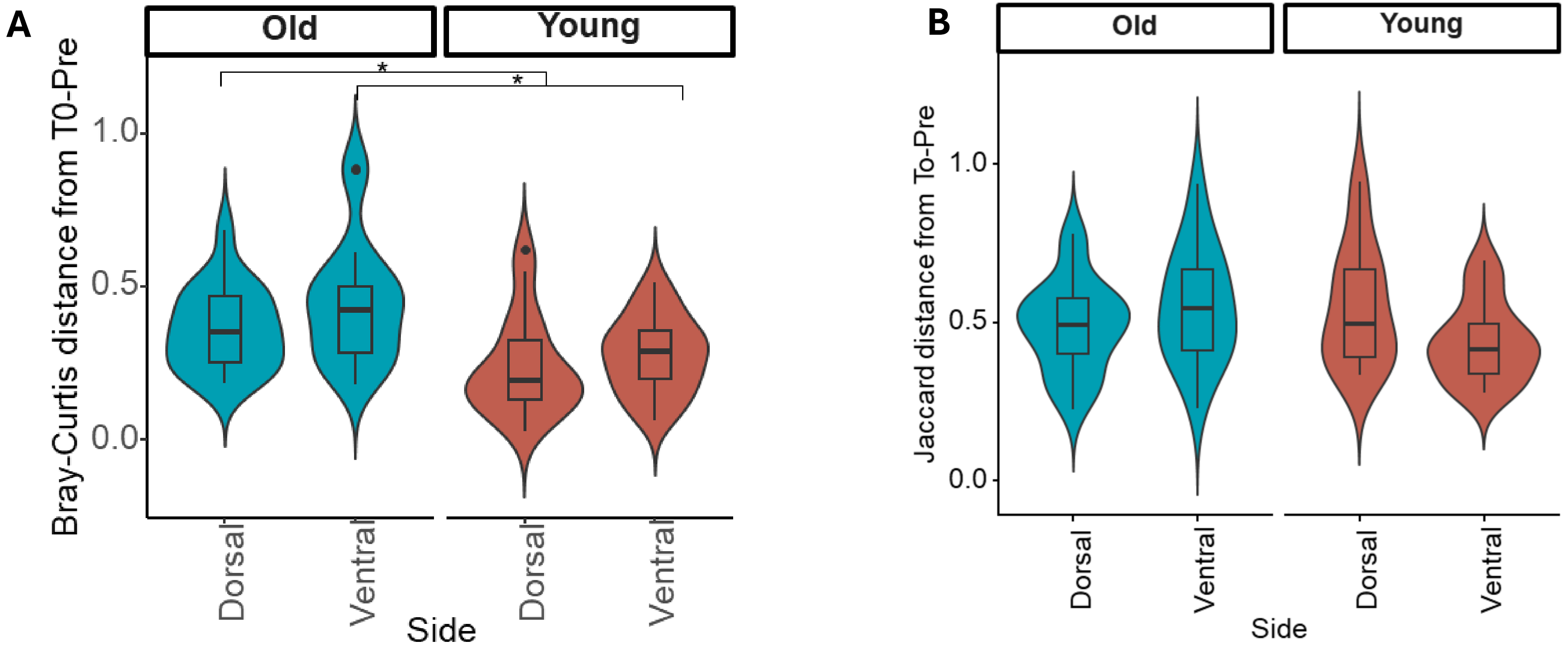


**Supplementary Figure 8 – Effect of tape stripping on microbial beta diversity:** Violin plots with inset boxplots showing the distribution of (A) Bray Curtis dissimilarity values and (B) Jaccard distance values between T0-Post and T0-Pre skin microbiomes. Paired Wilcoxon tests were used for within-group comparisons, and Wilcoxon rank-sum tests for between-group comparisons. False discovery rate (FDR)–adjusted p-values are denoted as “*” for $p \leq 0.05$. In all boxplots, the center line denotes the median, box limits the upper and lower quartiles, and whiskers the minimum and maximum values.

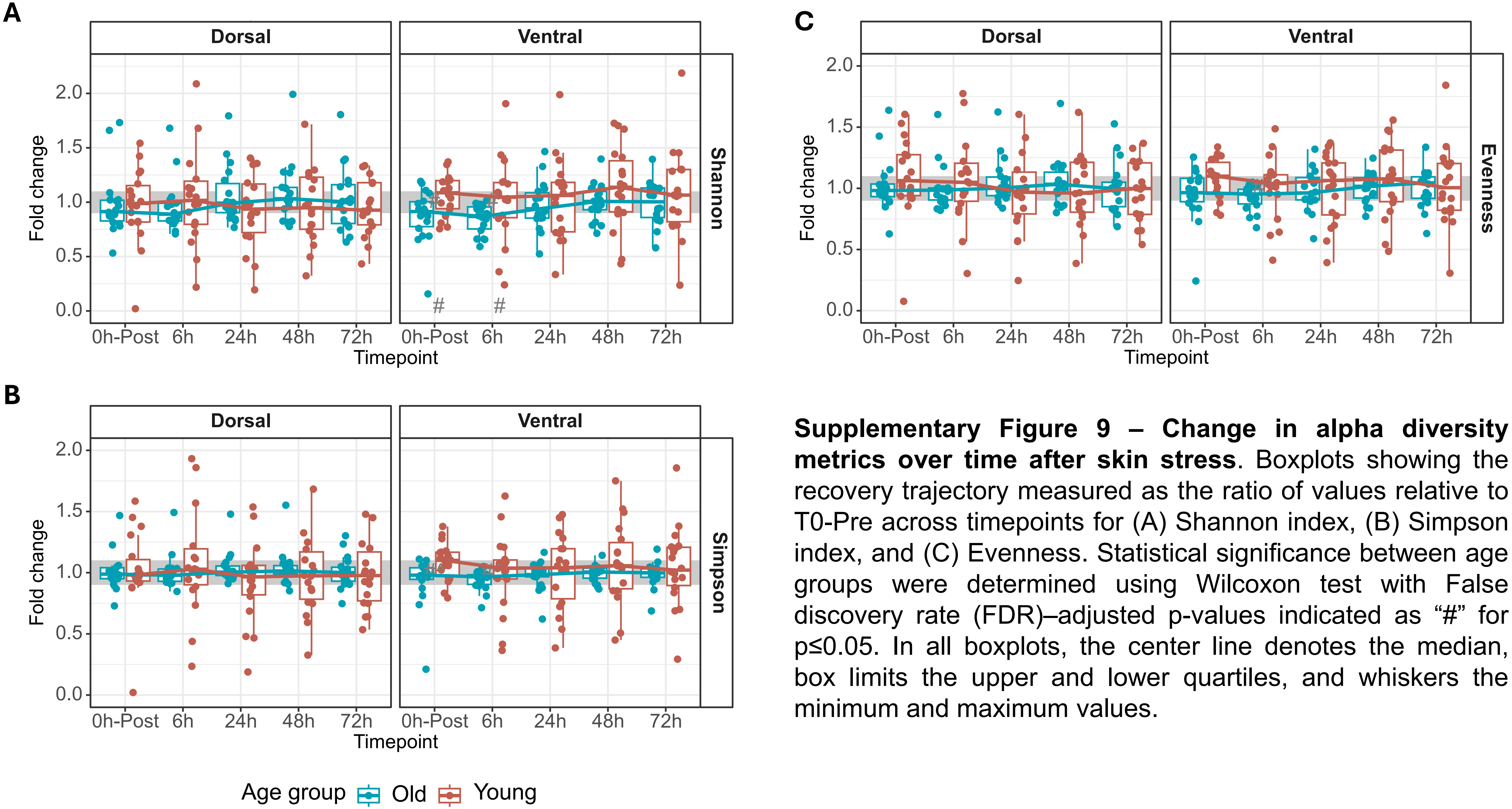


**Supplementary Figure 9 – Change in alpha diversity metrics over time after skin stress**. Boxplots showing the recovery trajectory measured as the ratio of values relative to T0-Pre across timepoints for (A) Shannon index, (B) Simpson index, and (C) Evenness. Statistical significance between age groups were determined using Wilcoxon test with False discovery rate (FDR)–adjusted p-values indicated as "#" for p≤0.05. In all boxplots, the center line denotes the median, box limits the upper and lower quartiles, and whiskers the minimum and maximum values.

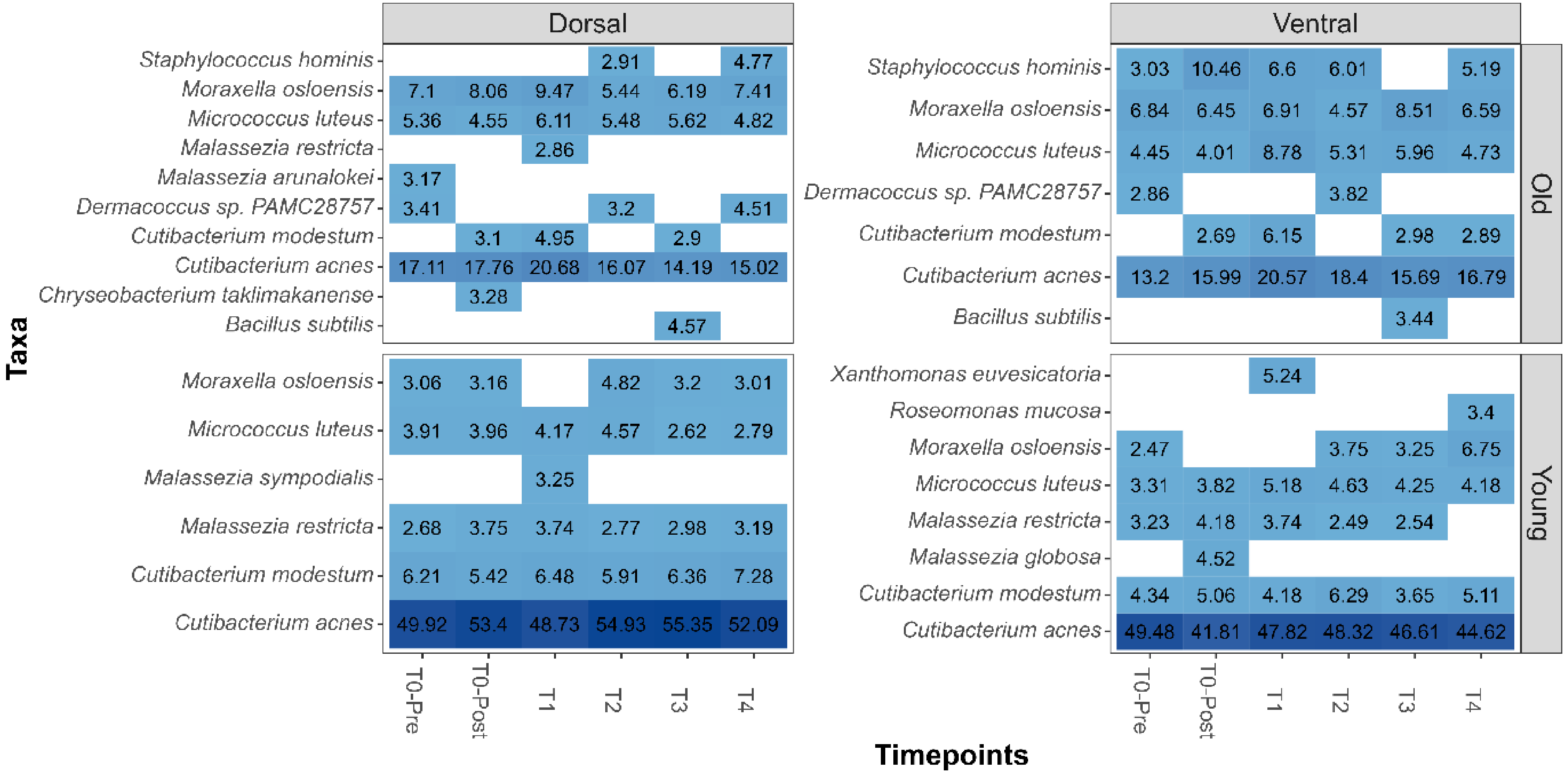


**Supplementary Figure 10 - Heatmap showing the mean variation of abundant species across timepoints.** Heatmap showing the variation across time of abundant species identified per group (top 5 selected per timepoint). Values and colours within the tile represent means.

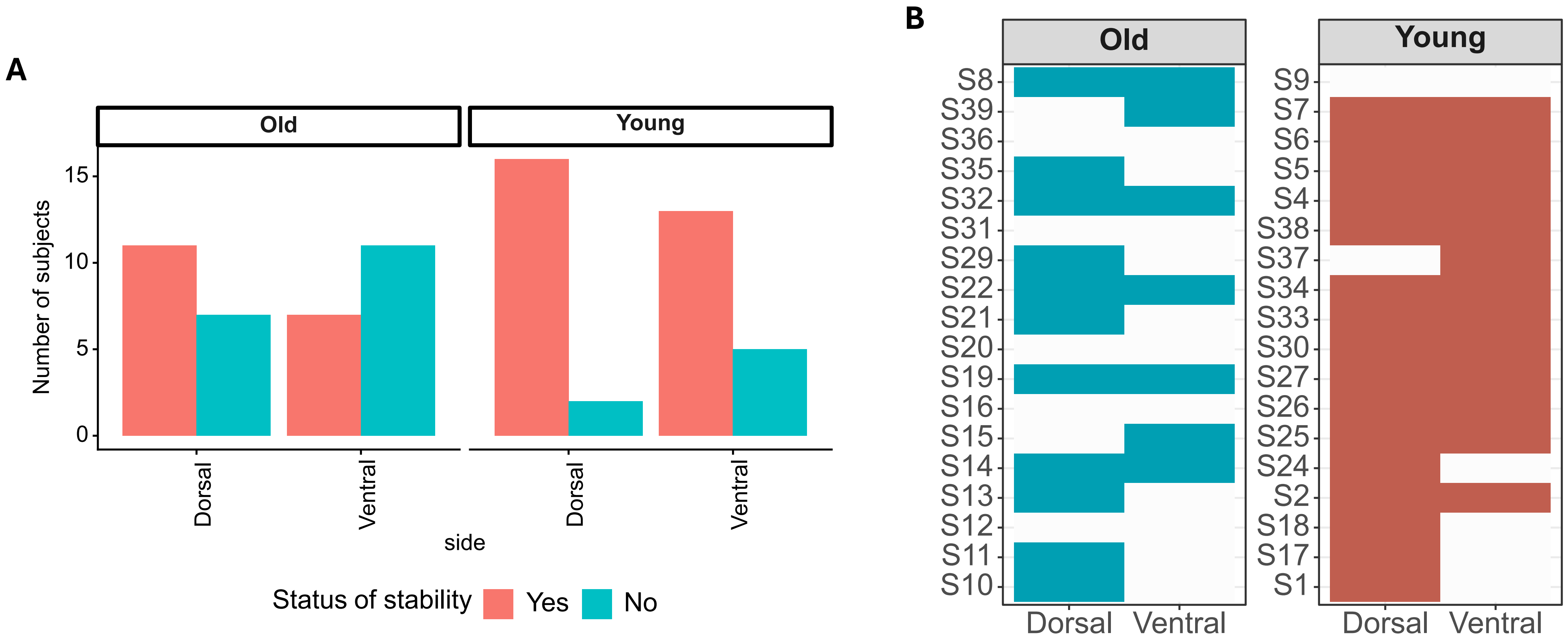


**Supplementary Figure 11 – Overall and individual microbial stability patterns (A)** Number of samples that attained microbial stability in each age group and site. Bars represent the count of subjects whose microbiome profiles reached a stable state based on the predefined stability criteria **(B)** Tile plot showing the attainment of microbiome stability (filled tile) across subjects in the old (left) and young (right) age groups, respectively, in dorsal and ventral sites.

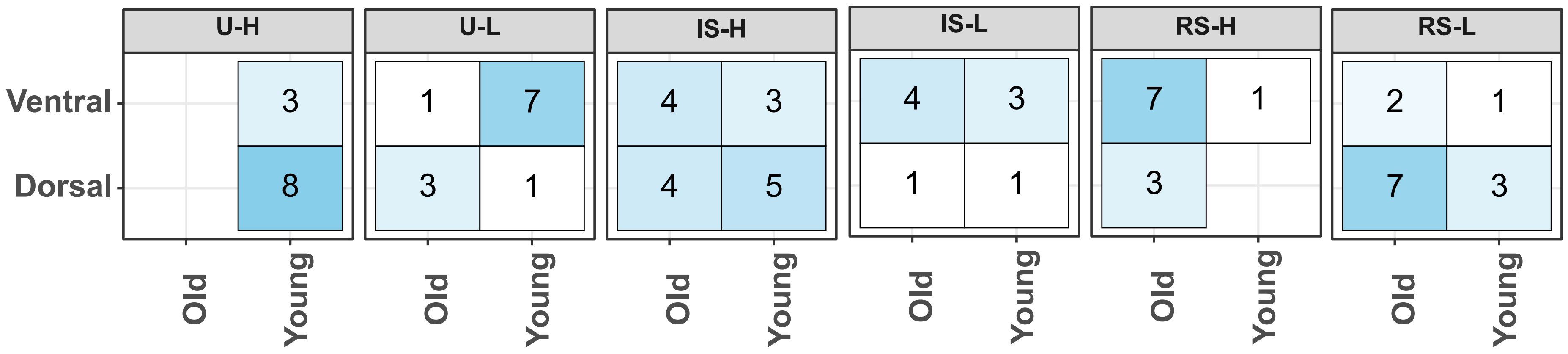


**Supplementary Figure 12 – Group constituents:** Heatmap showing the number of individuals in the microbial trajectory classes, stratified by age group and site.

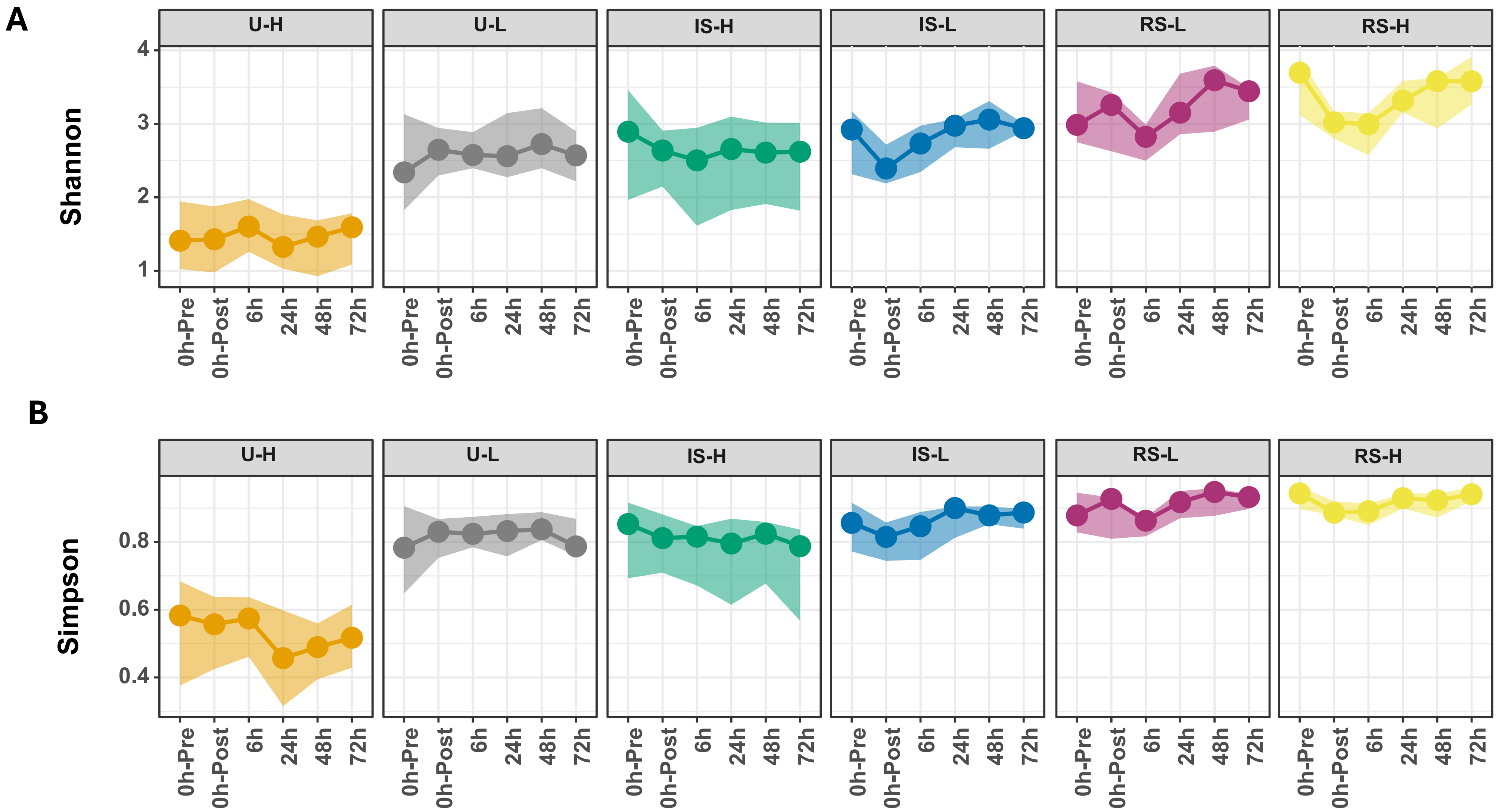


**Supplementary Figure 13 – Alpha diversity metrics across stability groups.** Line graph showing the temporal variation in (A) Shannon diversity, and (B) Simpson diversity across all six stability groups. The points represent the median values, and the shaded areas represent the interquartile range (first and third quartiles).

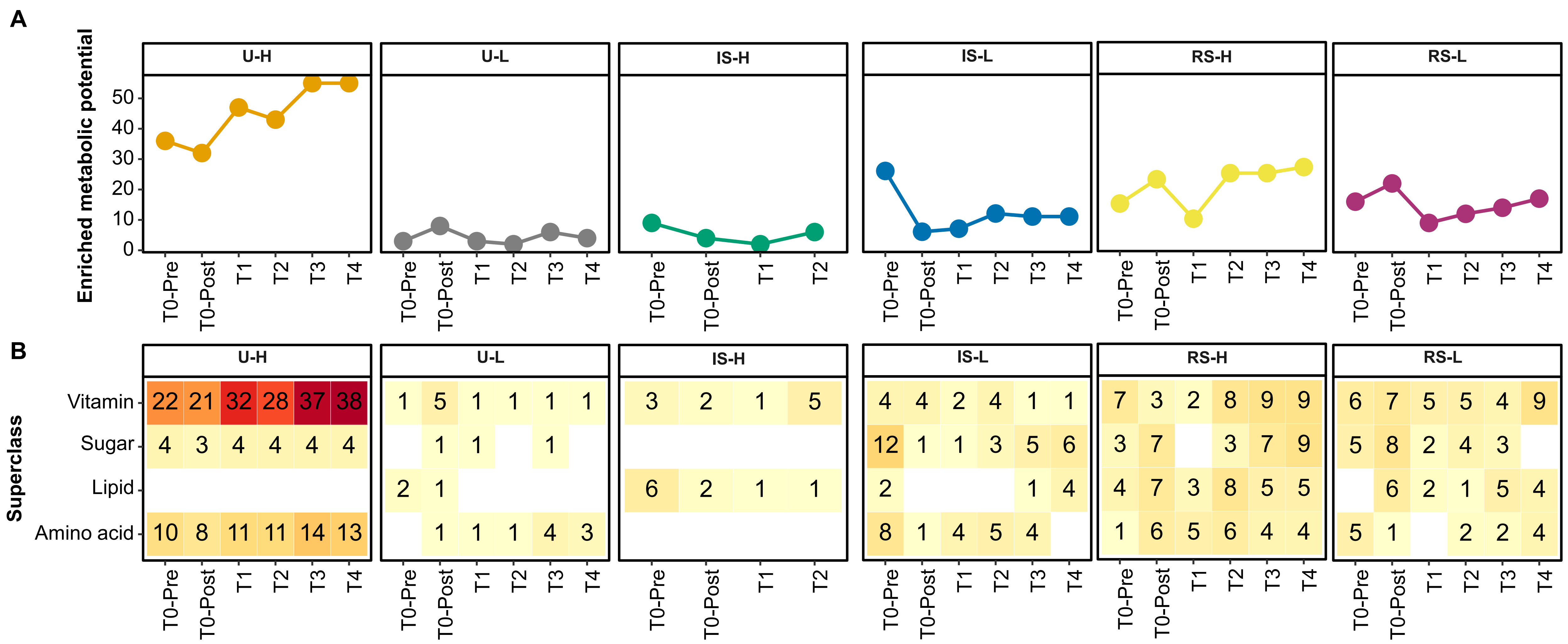


**Supplementary Figure 14 – Pathway enrichments across stability groups.** (A) Line plot showing the number of pathways enriched at each timepoint. (B) Heatmap showing the count of pathways within each subclass (y-axis) across various timepoints in each stability group.

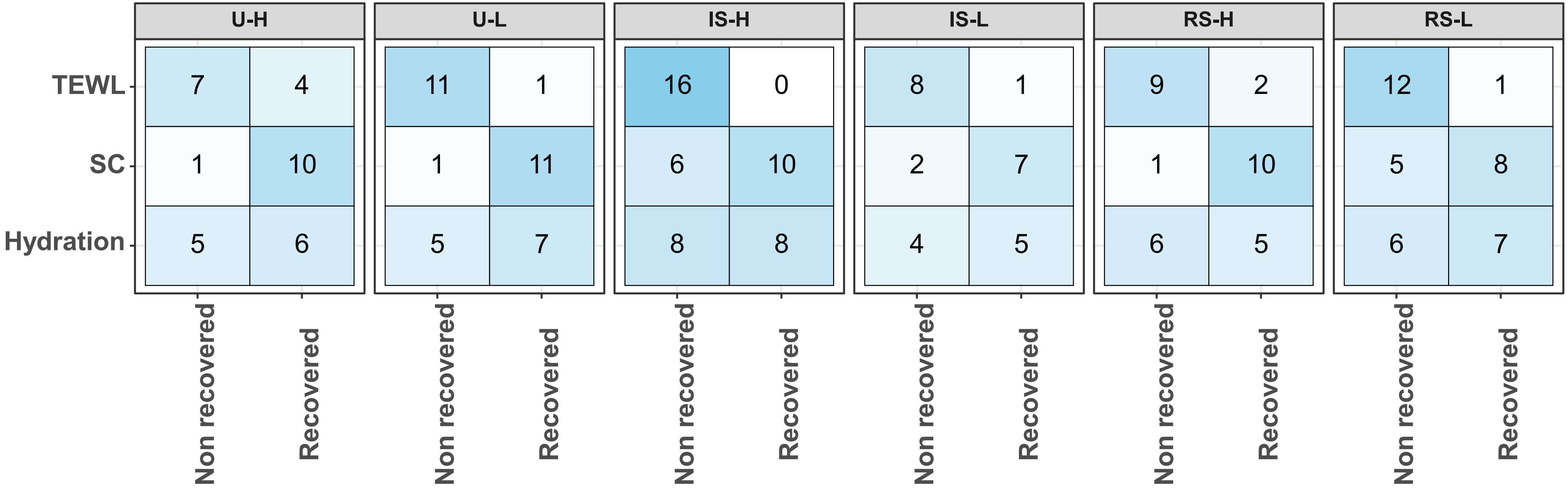


**Supplementary Figure 15 – Recovery statistics for skin physiological parameters within each stability group.** Heatmap showing the recovery statistics of members in the microbial trajectory classes.

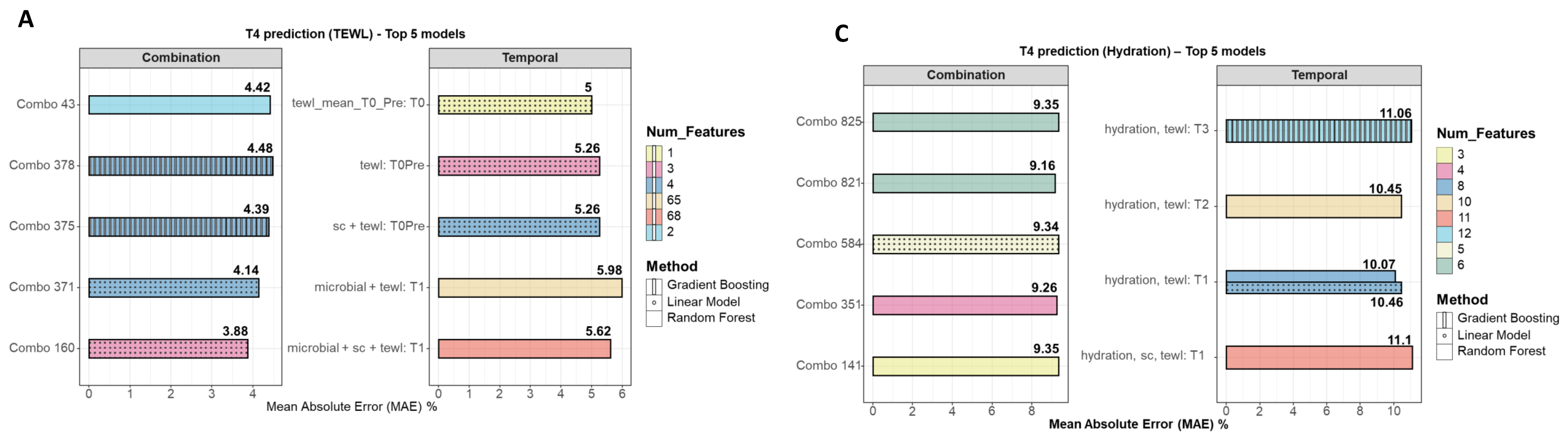


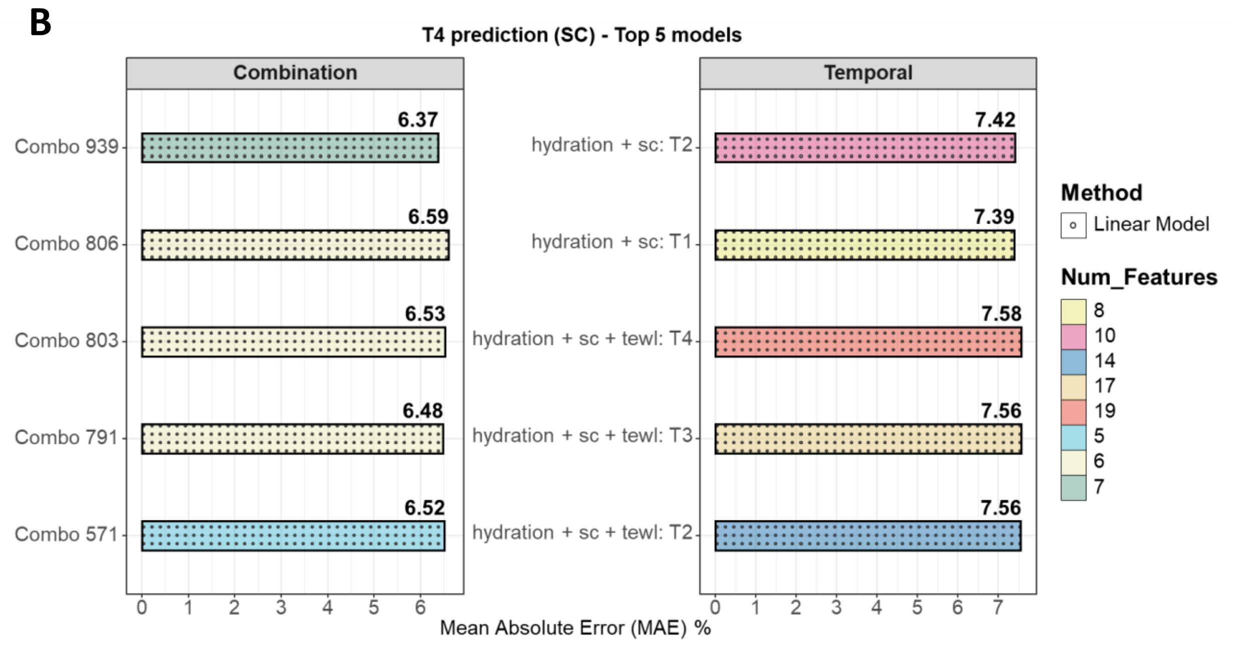


**Supplementary Figure 16 – Machine learning models to predict skin physiological parameters at 72 hours.** Bar plots showing the mean absolute error for 0-1 scaled values for the top 5 models under each category (Combination and Temporal – See Methods). (A) TEWL prediction at T4 (B) Stratum corneum thickness prediction at T4, and (C) Hydration prediction at T4. For the full list of features for every combination, see **Supplementary Data File 6**.

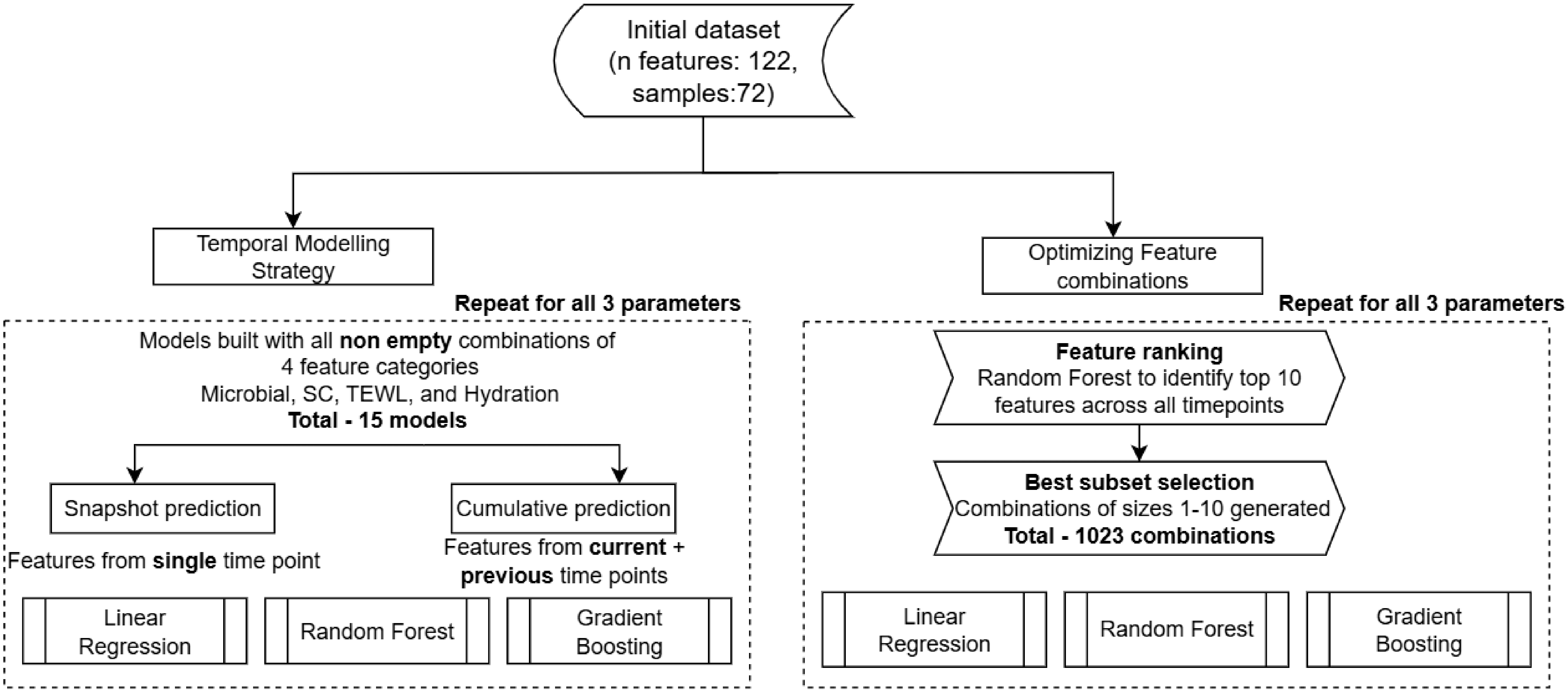


**Supplementary Figure 17 –** Flowchart showing the methodology used in training machine learning models to predict endpoint values
